# A mean-field toolbox for spiking neuronal network model analysis

**DOI:** 10.1101/2021.12.14.472584

**Authors:** Moritz Layer, Johanna Senk, Simon Essink, Alexander van Meegen, Hannah Bos, Moritz Helias

## Abstract

Mean-field theory of spiking neuronal networks has led to numerous advances in our analytical and intuitive understanding of the dynamics of neuronal network models during the past decades. But, the elaborate nature of many of the developed methods, as well as the difficulty of implementing them, may limit the wider neuroscientific community from taking maximal advantage of these tools. In order to make them more accessible, we implemented an extensible, easy-to-use open-source Python toolbox that collects a variety of mean-field methods for the widely used leaky integrate-and-fire neuron model. The Neuronal Network Mean-field Toolbox (NNMT) in its current state allows for estimating properties of large neuronal networks, such as firing rates, power spectra, and dynamical stability in mean-field and linear response approximation, without running simulations on high performance systems. In this article we describe how the toolbox is implemented, show how it is used to calculate neuronal network properties, and discuss different use-cases, such as extraction of network mechanisms, parameter space exploration, or hybrid modeling approaches. Although the initial version of the toolbox focuses on methods that are close to our own past and present research, its structure is designed to be open and extensible. It aims to provide a platform for collecting analytical methods for neuronal network model analysis and we discuss how interested scientists can share their own methods via this platform.

## 1 INTRODUCTION

Biological neuronal networks are composed of large numbers of recurrently connected neurons, with a single cortical neuron typically receiving synaptic inputs from thousands of other neurons (Braitenberg and Schüz, 1998; DeFelipe et al., 2002). Although the inputs of distinct neurons can be integrated in a complex fashion, average properties of entire populations of neurons often do not depend strongly on the contributions of individual neurons. Based on this observation, it is possible to develop analytically tractable theories of population properties, in which the effects of individual neurons are averaged out and the complex, recurrent input to individual neurons is replaced by a simplification, known as self-consistent effective input (Gerstner et al., 2014). In classical physics terms (e.g., Goldenfeld, 1992), this effective input is called *mean-field*, because it is the self-consistent mean of a *field*, which here is just another name for the input the neuron is receiving. The term *self-consistent* refers to the fact that the population of neurons that receives the effective input, is the same that contributes to this very input in a recurrent fashion: the population’s output determines its input and vice-versa. The stationary statistics of the effective input therefore can only be found in a self-consistent manner: the input to a neuron must be set exactly such that the caused output precisely leads to the respective input.

These mean-field theories have been developed for many different kinds of synapse, neuron, and network models. They have been successfully applied to study average population firing rates (van Vreeswijk and Sompolinsky, 1996; van Vreeswijk and Sompolinsky, 1998; Amit and Brunel, 1997b), and the various activity states a network of spiking neurons can exhibit, depending on the network parameters (Amit and Brunel, 1997a; Brunel, 2000; Ostojic, 2014), as well as the effects that different kinds of synapses have on firing rates (Fourcaud and Brunel, 2002; Lindner, 2004; Schuecker et al., 2015; Schwalger et al., 2015; Mattia et al., 2019). They have been used to investigate how neuronal networks respond to external inputs (Lindner and Schimansky-Geier, 2001; Lindner and Longtin, 2005), and they explain why neuronal networks can track external input on much faster time scales than a single neuron could (van Vreeswijk and Sompolinsky, 1996; van Vreeswijk and Sompolinsky, 1998). They allow studying pair-wise correlations of neuronal activity (Sejnowski, 1976; Ginzburg and Sompolinsky, 1994; Lindner et al., 2005; Trousdale et al., 2012) and were able to reveal why pairs of neurons in random networks, despite receiving a high proportion of common input, can show low output correlations (Hertz, 2010; Renart et al., 2010; Tetzlaff et al., 2012; Helias et al., 2014), which for example has important implication for information processing. They describe pair-wise correlations in network with spatial organization (Rosenbaum and Doiron, 2014; Rosenbaum et al., 2017; Dahmen et al., 2021) and can be generalized to correlations of higher orders (Buice and Chow, 2013). Mean-field theories were utilized to show that neuronal networks can exhibit chaotic dynamics (Sompolinsky et al., 1988; van Vreeswijk and Sompolinsky, 1996; van Vreeswijk and Sompolinsky, 1998), in which two slightly different initial states can lead to totally different network responses, which has been linked to the network’s memory capacity (Toyoizumi and Abbott, 2011; Schuecker et al., 2018). Most of the results mentioned above have been derived for networks of either rate, binary, or spiking neurons of a linear integrate-and-fire type. But various other models have been investigated with similar tools as well; for example, just to mention a few, Hawkes processes, non-linear integrate-and-fire neurons (Brunel and Latham, 2003; Fourcaud-Trocmé et al., 2003; Richardson, 2007, 2008; Grabska-Barwinska and Latham, 2014; Montbrió et al., 2015), or Kuramoto-type models (Stiller and Radons, 1998; van Meegen and Lindner, 2018). Additionally, there is an ongoing effort showing that many of the results derived for distinct models are indeed equivalent and that those models can be mapped to each other under certain circumstances (Grytskyy et al., 2013; Ostojic and Brunel, 2011; Senk et al., 2020).

Other theories for describing mean population rates in networks with spatially organized connectivity, based on taking a continuum limit, have been developed. These more phenomenological theories, known as neural field theories, have deepened our understanding of spatially and temporally structured activity patterns emerging in cortical networks. Starting with the seminal work by Wilson and Cowan (1972, 1973) investigating global activity patterns and Amari (1975, 1977) investigating stable localized neuronal activity. They were successfully applied to explain hallucination patterns (Ermentrout and Cowan, 1979; Bressloff et al., 2001), as well as EEG and MEG rhythms (Nunez, 1974; Jirsa and Haken, 1996, 1997). The neural field approach has been used to model working memory (Laing et al., 2002; Laing and Troy, 2003), motion perception (Giese, 2012), cognition (Schöner, 2008), and more; for extensive reviews of the literature, we refer the reader to Coombes (2005), Bressloff (2012), and Coombes et al. (2014).

Apparently, analytical theories have contributed to our understanding of neuronal networks and they provide a plethora of powerful and efficient methods for network model analysis. However, following the details behind those theories often requires some mathematical background. Furthermore, the analytical methods developed in those works only are of immediate use to other researchers if there is a numerical implementation that is flexible enough to be applicable to different network models, according to that researchers’ questions. Such an implementation is often far from straightforward and at times requires investing substantial time and effort. Commonly, such tools are implemented as the need arises, and their reuse is often not organized systematically. Also reuse is often restricted to within a single lab. This way, not only are effort and costs spent by the neuroscientific community duplicated over and over again, but also are many scientists deterred from taking maximal advantage of those methods; simply because they do not have the capacities to do so, although such tools might open new avenues for investigating their research questions.

In order to make analytical tools for neuronal network model analysis accessible to a wider part of the neuroscientific community, and to create a platform for collecting well-tested and validated implementations of such tools, we have developed the Python toolbox NNMT (Layer et al., 2021), short for Neuronal Network Mean-field Toolbox. NNMT has been designed to fit the diversity of mean-field theories, and the key features we are aiming for are modularity, extensibility, and a simple usability. Furthermore, it features an extensive test suite to ensure the validity of the implementations as well as a comprehensive user documentation. The current version of NNMT mainly comprises tools for investigating networks of leaky integrate-and-fire neurons as well as some methods for studying neural field models. The toolbox is open-source and publicly available on GitHub^1^.

There exist different approaches for neuronal network model analysis that do not use direct simulations of the subthreshold dynamics of individual neurons. Cain et al. (2016), for example, analyze the Potjans and Diesmann cortical microcircuit model (Potjans and Diesmann, 2014) employing the simulation platform DiPDE^2^. DiPDE is based on the population density approach and yields an efficient numerical method for analyzing the statistical evolution of voltage distributions of large homogeneous populations of neurons. Schwalger et al. (2017) start from a stochastic microscopic neuron model and derive mesoscopic population equations, which reproduce the statistical and qualitative behavior of the homogeneous neuronal sub-populations. But to the best of our knowledge, there currently is no toolbox pursuing efforts to collect numerical implementations of various analytical mean-field tools for neuronal network model analysis.

**Listing 1:**
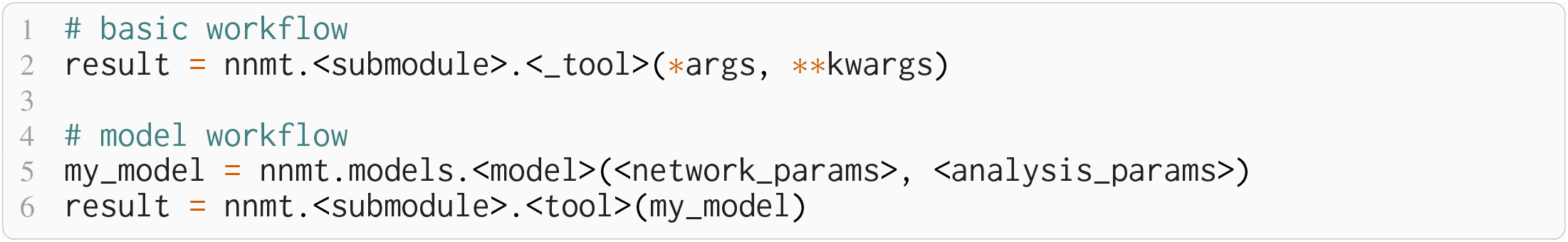
The two modes of using NNMT: In the basic workflow (top), quantities are calculated by passing all required arguments directly to the underscored tool functions available in the submodules of NNMT. In the model workflow (bottom), a model class is instantiated with parameter sets and the model instance is passed to the non-underscored tool functions which automatically extract the relevant parameters.

In the following, we present the design considerations that led to the structure and implementation of NNMT as well as a representative set of use cases. Section 2 first introduces its architecture. Section 3 then explains its usage by reproducing previously published network model analyses from Sanzeni et al. (2020), Bos et al. (2016), Schuecker et al. (2015), and Senk et al. (2020). Section 4 discusses the role of community-wide accessible toolboxes for scientific software development in general and indicates current limitations and prospective advancements of NNMT.

## 2 WORKFLOWS AND ARCHITECTURE

What are the requirements a package for collecting analytical methods for neuronal network model analysis needs to fulfill? To begin with, it must be adaptable and modular enough to accommodate many and diverse analytical methods while avoiding code repetition and a complex interdependency of package components. It must enable the application of the collected algorithms to various network models in a simple and transparent manner. It needs to make the tools easy to use for new users, while also providing experts with direct access to all parameters and options. Finally, the methods need to be thoroughly tested and well documented.

These are the main considerations that guided the development of NNMT. **Figure 1A** illustrates that the toolbox can be used in to two different workflows depending on the preferences and goals of the user. In the *basic workflow* the individual method implementations called *tools* are directly accessed, whereas the *model workflow* provides additional functionality for the handling of parameters and results.

**Figure 1.**
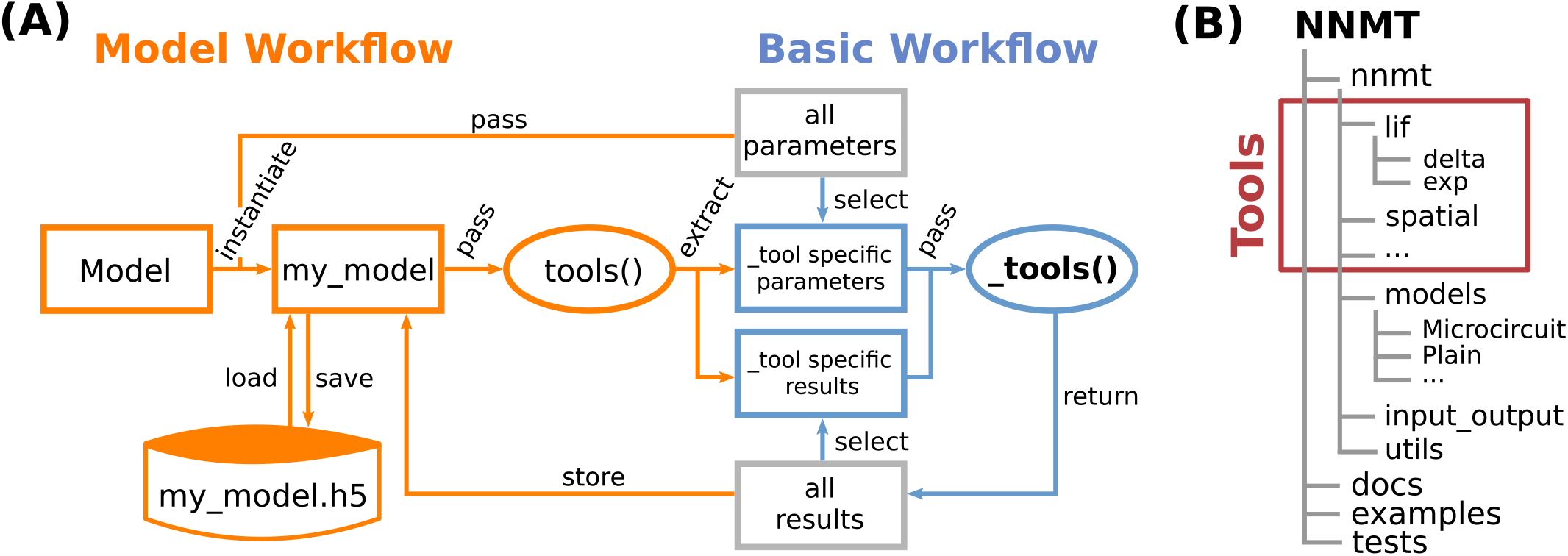
The Neuronal Network Mean-field Toolbox (NNMT) is a platform for collecting network analysis tools. **(A)** Individual tool functions can be used directly by explicitly passing arguments (basic workflow, blue) or via a convenience layer with a model class and wrapper functions facilitating the handling of parameters and results (model workflow, orange). **(B)** Structure of the repository. In addition to the tool collection (red frame, containing the `tools()` and the `_tools())`, pre-defined model classes, and utility functions in the NNMT Python package, the repository also comprises documentation, usage examples, and a test suite.

### 2.1 Basic Workflow

The core of NNMT is a collection of low-level functions that take specific parameters (or pre-computed results) as input arguments and return analytical results of network properties. In **Figure 1A**, we refer to such basic functions as `_tools()`, beginning with an underscore. We term this lightweight approach of directly using these functions the basic workflow. The top part of **Listing 1** demonstrates this usage; for example, the quantity to be computed could be the mean firing rate of a neuronal population and the arguments could be parameters which define neuron model and external drive. While the basic workflow gives full flexibility and direct access to every parameter of the calculation, it remains the user’s responsibility to insert the arguments correctly, e.g., in the right units.

### 2.2 Model Workflow

The model workflow adds a convenience layer on top of the basic workflow **(Figure 1A**). A *model* in this context is an object that stores a larger set of parameters and can be passed directly to a `tool()`, the non-underscored wrapper of the respective `_tool()`. The `tool()` automatically extracts the relevant parameters from the model, passes them as arguments to the corresponding core function `_tool()`, returns the results, and stores them in the model. The bottom part of **Listing 1** shows how a model is initialized with parameters and then passed to a `tool()` function.

Models are implemented as Python classes and can be found in the submodule `nnmt.models`. We provide the class `nnmt.models.Network` as a parent class and a few example child classes which inherit the generic methods and properties but are tailored to specific network models; custom models can be added straightforwardly. The parameters distinguish network parameters, which define neuron models and network connectivity, and analysis parameters; an example for an analysis parameter is a frequency range over which a function is evaluated. Upon model instantiation, parameter sets defining values and corresponding units are passed as Python dictionaries or `yaml` files. The model constructor takes care of reading in these parameters, deriving dependent parameters, and converting all units to SI units for internal computations. Consequently, the parameters passed as arguments and the functions for deriving dependent parameters of a specific child class need to be aligned. This design encourages a clear separation between a concise set of base parameters and functionality that transforms these parameters to the generic (vectorized) format that the tools work with. To illustrate this, consider the weight matrix of a network of excitatory and inhibitory neuron populations in which all excitatory connections have the same weight and all inhibitory ones another weight. As argument one could pass just a tuple of these two different weight values and the corresponding model class would take care of constructing the full weight matrix.

When a `tool()` is called, the function will check whether all required parameters are available in the passed model object. If not, an error is raised, pointing the user towards the missing parameters. If the property to be computed is based on the computation of another property that has yet to be calculated, the `tool()` will also raise an error, informing the user of which other properties must be calculated first. This way, the user is guided towards the desired result. When all required parameters and results are available, the `tool()` extracts the required arguments, calls the respective `_tool()`, caches the result and its units into the network model’s result dictionaries, and returns the result. If the user attempts to compute the same property twice, using identical parameters, the `tool()` will retrieve the already computed result from the cache and return that value. Results can be exported to an HDF5 file and also loaded.

Using the model workflow instead of the basic workflow comes with the initial overhead of choosing a suitable combination of parameters and a model class, but has the advantages of a higher level of automation with built-in mechanisms for checking correctness of input (e.g., regarding units), reduced redundancy, and the options to store and load results. Both modes of using the toolbox can also be combined.

### 2.3 Structure of the Toolbox

The structure of the NNMT repository is depicted in **Figure 1B**. The Python package `nnmt` is sub-divided into submodules containing the model classes (`nnmt.models`), helper routines (at present `nnmt.input_output` and `nnmt.utils`), and the tools. The tools are organized in a modular, extensible fashion with a streamlined hierarchy. To give an example, a large part of the currently implemented tools apply to networks of leaky integrate-and-fire (LIF) neurons and they are located in the submodule `nnmt.lif`. The mean-field theory for networks of LIF neurons distinguishes between neurons with instantaneous synapses, also called delta synapses, and those with exponentially decaying post-synaptic currents. Similarly, the submodule for LIF neurons is split further into the two submodules `nnmt.lif.delta` and `nnmt.lif.exp`. NNMT also collects different implementations for computing the same quantity using different approximations or numerics, allowing for a comparison of different approaches.

Apart from the core package, NNMT comes with an extensive online documentation^3^, including a quickstart tutorial, all examples presented in this paper, a complete documentation of all tools, as well as a guide for contributors.

Furthermore, we provide an extensive test suite that validates the tools by checking them against previously published results and alternative implementations where possible. This ensures that future improvements of the numerics do not break the tools.

## 3 HOW TO USE THE TOOLBOX

In this section, we demonstrate the practical use of NNMT by replicating a variety of previously published results. The examples presented have been chosen to cover a broad range of common use cases and network models. We include analyses of both stationary and dynamic network features, as mean-field theory is typically divided into two parts: stationary theory, which describes time-independent network properties of systems in a stationary state, and dynamical theory, which describes time-dependent network properties. Additionally, we show how to use the toolbox to map a spiking to a simpler rate model, as well as how to perform a linear stability analysis.

### 3.1 Installation and Setup

The toolbox can be either installed using pip

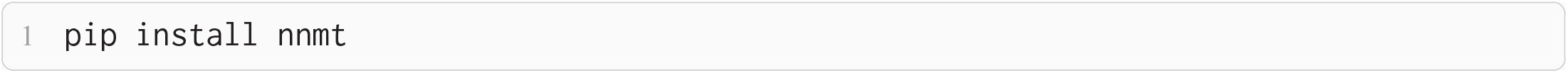

or by installing it directly from the repository, which is described in detail in the online documentation. After the installation, the module can be imported:

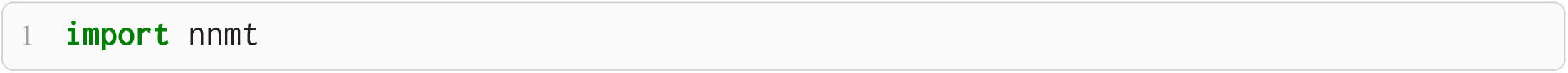

### 3.2 Stationary Quantities

#### 3.2.1 Response Nonlinearities

Networks of excitatory and inhibitory neurons **(Figure 2A**) are widely used in computational neuroscience (Gerstner et al., 2014), e.g., to show analytically that a balanced state featuring asynchronous, irregular activity emerges dynamically in a broad region of the parameter space (Brunel, 2000). How do these networks respond to external input? Sanzeni et al. (2020) uncover five different types of nonlinearities in the network response depending on the network parameters. Here, we show how to use the toolbox to reproduce their result **(Figure 2B–F**).

**Figure 2.**
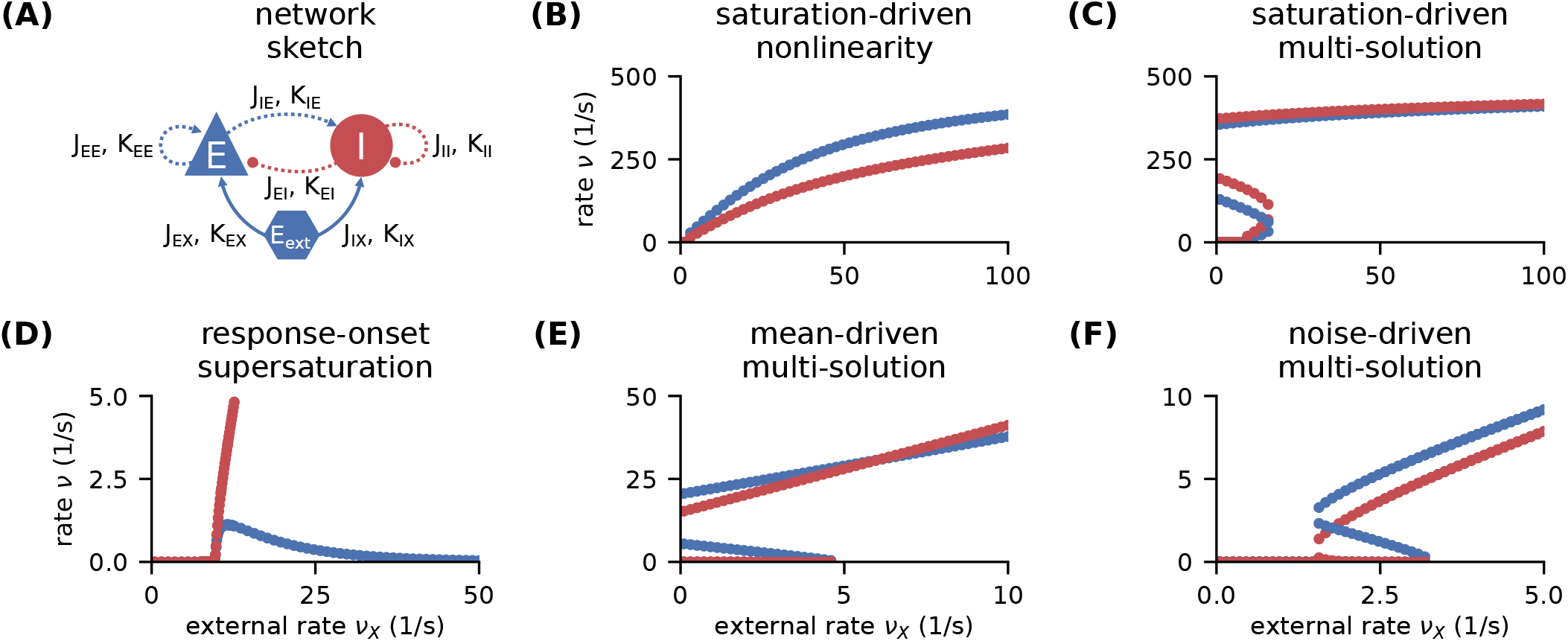
Response nonlinearities in EI-networks. **(A)** Network diagram with nodes and edges according to the graphical notation proposed by Senk et al. (2021). **(B–F)** Firing rate of excitatory (blue) and inhibitory (red) population for varying external input rate *ν_X_*. Specific choices for synaptic weights (***J***, ***J***_ext_) and in-degrees (***K***, ***K***_ext_) lead to five types of nonlinearities: **(B)** saturation-driven nonlinearity, **(C)** saturation-driven multi-solution, **(D)** response-onset supersaturation, **(E)** mean-driven multi-solution, and **(F)** noise-driven multi-solution. See Figure 8 in (Sanzeni et al., 2020) for parameters.

The network consists of two populations of identical LIF neurons with instantaneous (*delta*) synapses. The synaptic weights and mean in-degrees of recurrent and external connections are population-specific:

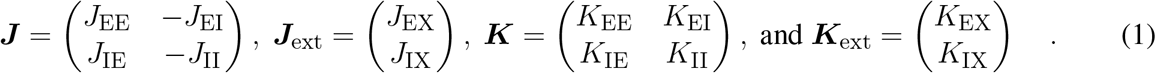

All weights are positive, implying an excitatory external input. Approximating the input to a neuron as Gaussian white noise *ξ*(*t*) with mean 〈*ξ*(*t*)〉 = *μ* and noise intensity 〈*ξ*(*t*)*ξ*(*t*′)〉 = *τ*_m_*σ*^2^*δ*(*t* – *t*′), its firing rate is given by (Siegert, 1951; Amit and Brunel, 1997b)

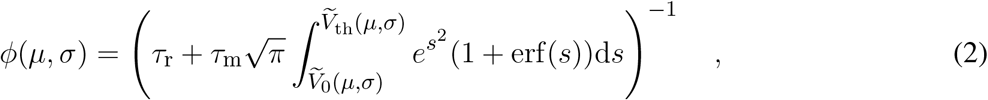

where *τ*_m_ denotes the membrane time constant, *τ*_r_ the refractory period, and the rescaled reset- and threshold-voltages are

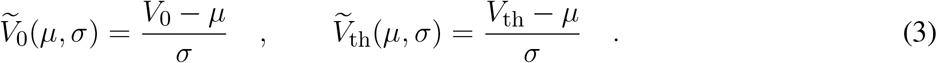

The first term in Eq. (2) is the refractory period and the second term is the mean first-passage time of the membrane voltage from reset to threshold. The mean and the noise intensity of the input to a neuron in a population *A*, which control the mean first-passage time through Eq. (3), are determined by (Amit and Brunel, 1997b)

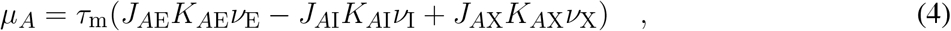

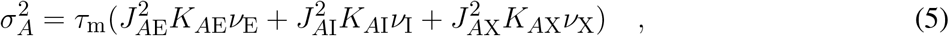

respectively, where each term reflects the contribution of one population, with the corresponding firing rates of the excitatory *ν*_E_, inhibitory *ν*_I_, and external population *ν*_X_. Note that the excitatory and inhibitory inputs counteract each other in *μ_A_*, Eq. (4), while their contributions add up in *σ_A_*, Eq. (5). Both *μ_A_* and *σ_A_* depend on the firing rate of the neurons *ν_A_*, which is in turn given by Eq. (2). Thus, one arrives at the self-consistency problem

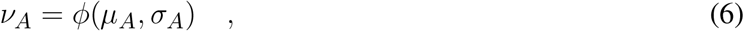

which is coupled across the populations due to Eq. (4) and Eq. (5).

Our toolbox provides two algorithms to solve Eq. (6): 1) Integrating the auxiliary ordinary differential equation (ODE) 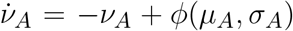 with initial values *ν_A_*(0) = *ν*_*A*,0_ until it reaches a fixed point 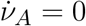, where Eq. (6) holds by construction. 2) Minimizing the quadratic deviation 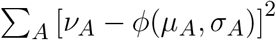, using least squares (LSTSQ) starting from an initial guess *ν*_*A*,0_. The ODE method is robust to changes in the initial values and hence a good first choice. However, it cannot find self-consistent solutions that correspond to an unstable fixed point of the auxiliary ODE. To this end, the LSTSQ method can be used. Its drawback is that it needs a good initial guess, because otherwise the found minimum might be a local one where the quadratic deviation does not vanish and which does not correspond to a self-consistent solution. A prerequisite for both methods is a numerical solution of the integral in Eq. (2); this is discussed in the Appendix, Section 5.1.

The solutions of the self-consistency problem Eq. (6) for varying *ν*_X_ and fixed ***J***, ***J***_ext_, ***K***, and ***K***_ext_ reveal the five types of response nonlinearities (**Figure 2**). Different response nonlinearities arise through specific choices of synaptic weights, **J** and ***J***_ext_, and in-degrees, ***K*** and ***K***_ext_, which suggests that already a simple EI-network possesses a rich capacity for nonlinear computations. Whenever possible, we use the ODE method and resort to the LSTSQ method only if the self-consistent solution corresponds to an unstable fixed point of the auxiliary ODE. Combining both methods, we can reproduce the first columns of Figure 8 in Sanzeni et al. (2020), where all five types of nonlinearities are presented. Curiously, some of the solutions corresponding to an unstable fixed point are nonetheless found in network simulations according to other panels of Figure 8 in Sanzeni et al. (2020) which we do not reproduce here.

In **Listing 2**, we show a minimal example to produce the data shown in **Figure 2B**. After importing the function that solves the self-consistency Eq. (6), we collect the neuron and network parameters in a dictionary. Then, we loop through different values for the external rate *ν_X_* and determine the network rates using the ODE method, which is sufficient in this example. In **Listing 2** and to produce **Figure 2**, we use the basic workflow because only one isolated tool of NNMT (`nnmt.lif.delta._firing_rates()`) is employed, which requires only a few parameters defining the simple EI-network.

**Listing 2:**
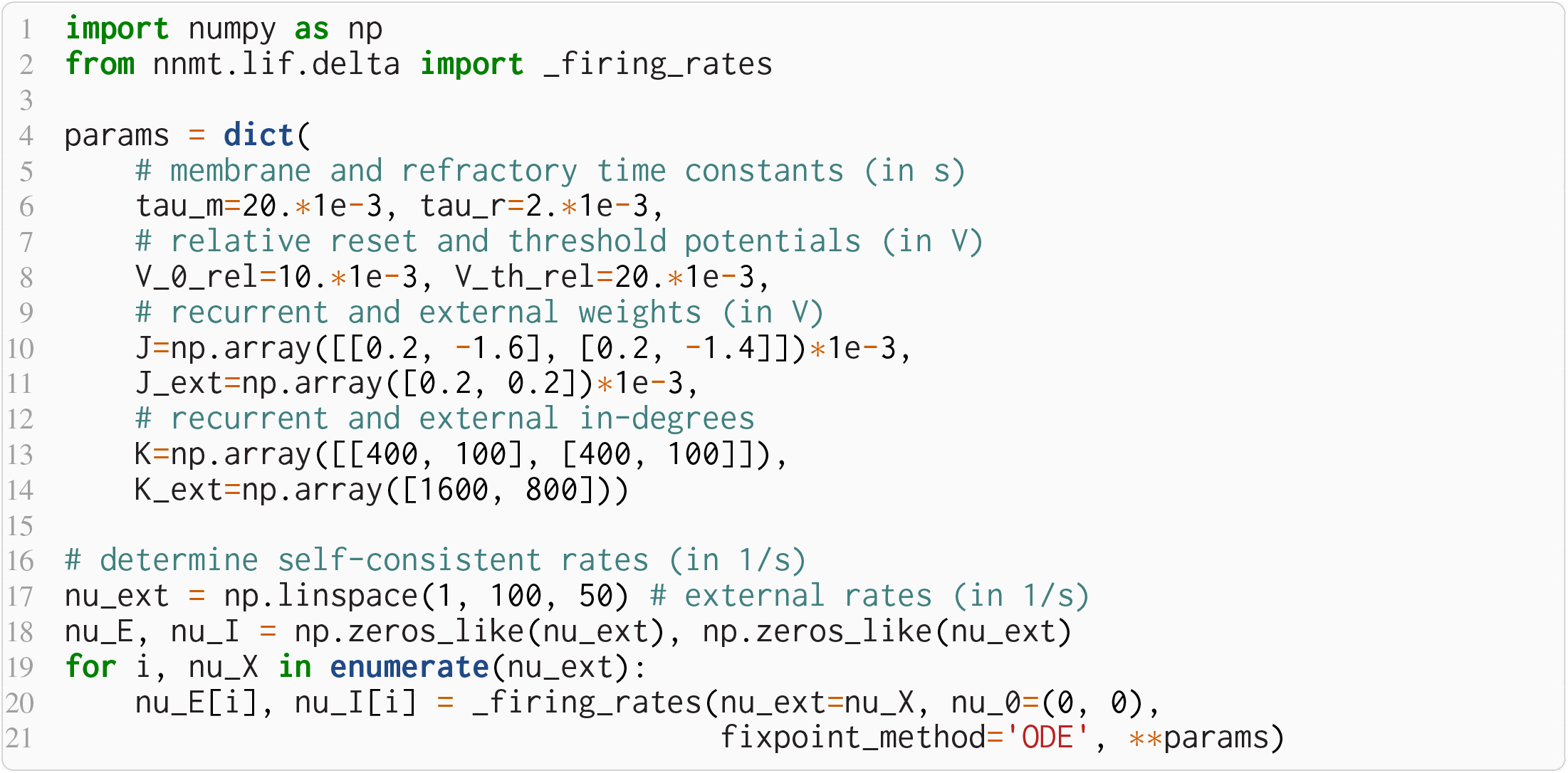
Example script to produce the data shown in **Figure 2B** using the ODE method (initial value *ν*_*A*,0_ = 0 for population *A* ∈ {*E*, *I*}).

#### 3.2.2 Firing Rates of Microcircuit Model

Here we show how to use the model workflow to calculate the firing rates of the cortical microcircuit model by Potjans and Diesmann (2014). The circuit is a simplified point neuron network model with biologically plausible parameters which has been recently used in a number of other works, for example to study network properties, to assess the performance of different simulators, and to extend it to large-scale models (e.g., Wagatsuma et al., 2011; Bos et al., 2016; Hagen et al., 2016; Schmidt et al., 2018; van Albada et al., 2018; Knight and Nowotny, 2018; Golosio et al., 2021; Dasbach et al., 2021). The model consists of eight populations of LIF neurons, corresponding to the excitatory and inhibitory populations of four cortical layers: 2/3E, 2/3I, 4E, 4I, 5E, 5I, 6E, and 6I (see **Figure 3A**). It defines the number of neurons in each population, the number of connections between the populations, the single neuron properties, and the external input. Simulations show that the model yields realistic firing rates for the different populations (Potjans and Diesmann, 2014).

**Figure 3.**
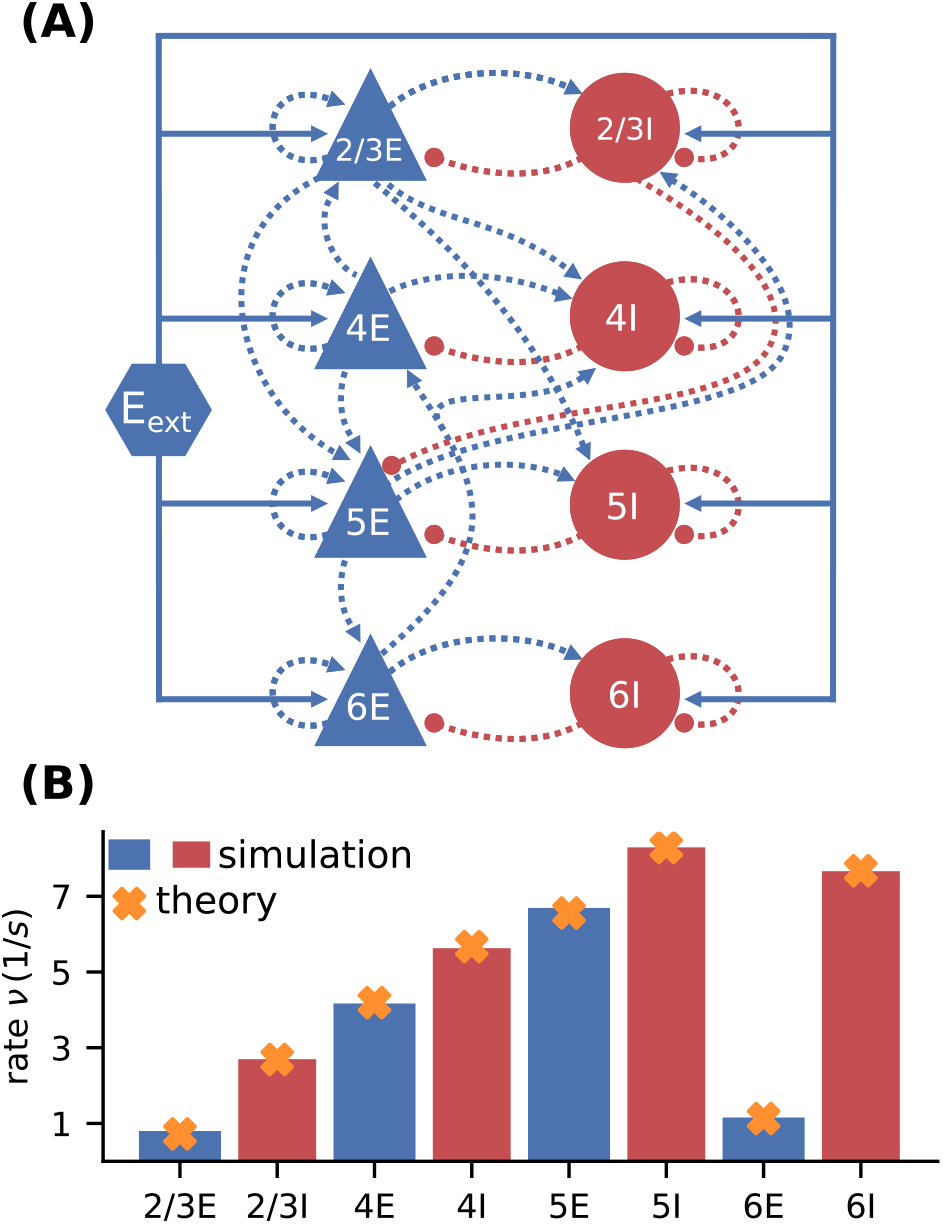
Cortical microcircuit model by Potjans and Diesmann (2014). **(A)** Network diagram (only the strongest connections are shown as in Figure 1 of the original publication). Same notation as in **Figure 2A**. **(B)** Simulation and mean-field estimate for average population firing rates using the parameters from Bos et al. (2016).

In contrast to the EI-network model investigated in Section 3.2.1, the neurons in the microcircuit model have exponentially shaped post-synaptic currents. Fourcaud and Brunel (2002) developed a method for calculating the firing rate for this synapse type. They have shown that, if the synaptic time constant *τ*_s_ is much smaller than the membrane time constant *τ*_m_, the firing rate for LIF neurons with exponential synapses can be calculated using Eq. (2) with shifted integration boundaries

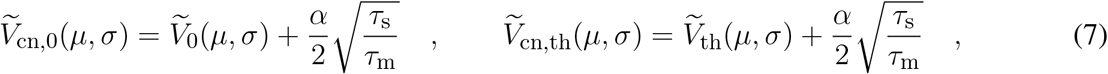

with the rescaled reset- and threshold-voltages from Eq. (3) and 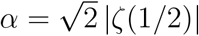, where *ζ*(*x*) denotes the Riemann zeta function; the subscript cn stands for “colored noise”.

The microcircuit has been implemented as an NNMT model (`nnmt.models.Microcircuit`). We here use the parameters of the circuit as published in Bos et al. (2016) which is slightly differently parameterized than the original model (see Appendix, **Table 1**). The model parameter values and units are defined using a `yaml` file, which uses a dictionary style syntax with `key:value` pairs:

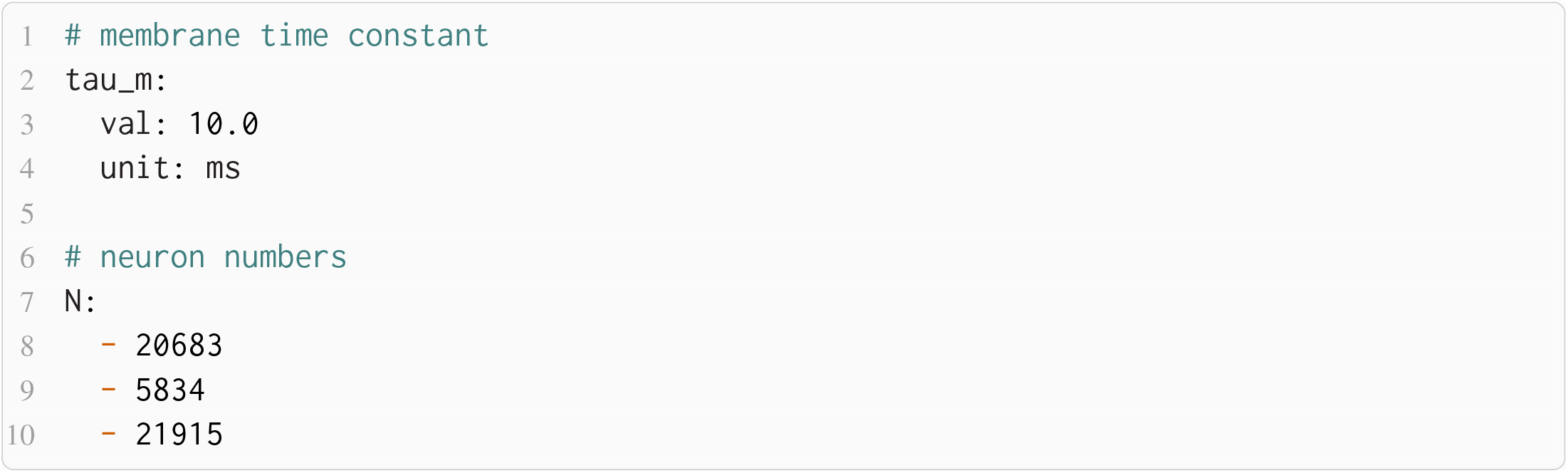

Once the parameters are defined, a microcircuit model is instantiated by passing the respective parameter file to the model constructor; the units are automatically converted to SI units. Then the firing rates are computed. For comparison, we finally load the simulated rates from Bos et al. (2016):

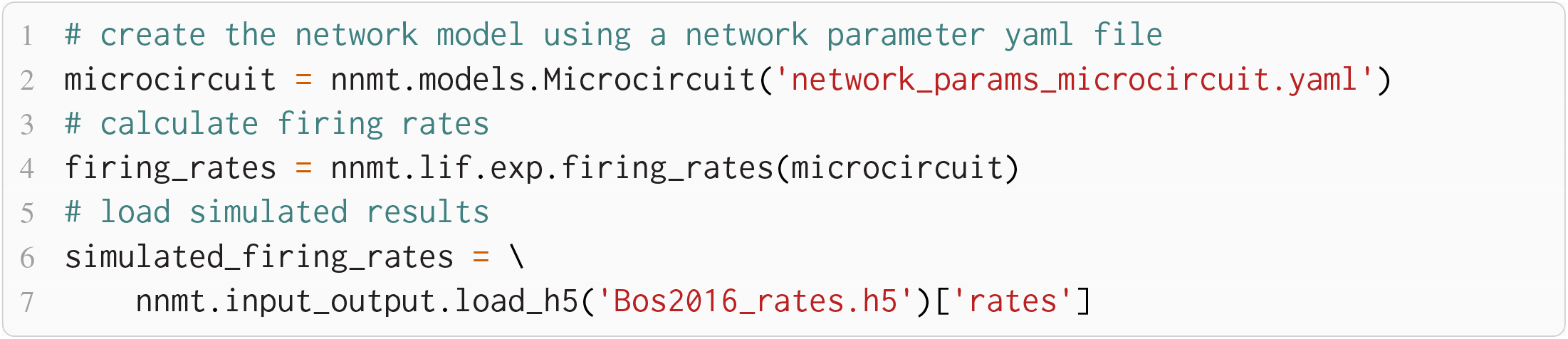

The simulated rates have been obtained by a numerical network simulation in which the neuron populations are connected according to the model’s original connectivity rule: random, fixed total number with multapses (autapses prohibited), see Senk et al. (2021). For simplicity, the theoretical predictions assume a connectivity with a fixed in-degree for each neuron. Dasbach et al. (2021) show that simulated spike activity data of networks with these two different connectivity rules are characterized by differently shaped rate distributions (“reference” in their Figures 3d and 4d). In addition, the weights in the simulation are normally distributed while the theory replaces each distribution by its mean; this corresponds to the case *N*_bins_ = 1 in Dasbach et al. (2021). Nevertheless, our mean-field theoretical estimate of the average population firing rates is in good agreement with the simulated rates **(Figure 3B**).

**Figure 4.**
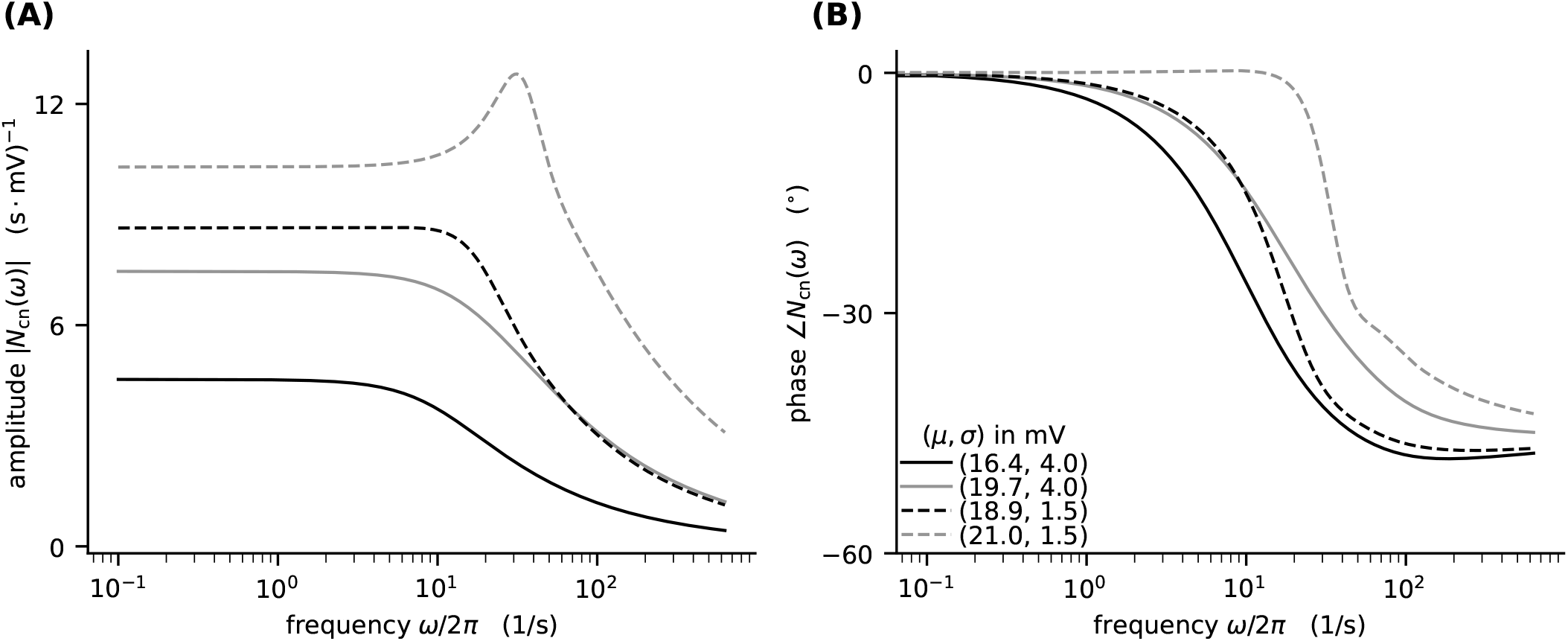
Colored-noise transfer function *N_cn_* of LIF model in different regimes. **(A)** Absolute value and **(B)** phase of the “shift” version of the transfer function as a function of the log-scaled frequency. Neuron parameters are set to *V*_th_ = 20 mV, *V*_0_ = 15 mv, *τ*_m_ = 20 ms, and *τ*_s_ = 0.5 ms. For given noise intensities of input current, *σ* = 4 mV (solid line) and *σ* = 1.5 mV (dashed line), the mean input *μ* is chosen such that firing rates *ν* = 10 Hz (black) and *ν* = 30 Hz (gray) are obtained.

### 3.3 Dynamical Quantities

#### 3.3.1 Transfer Function

One of the most important dynamical properties of a neuronal network is how it reacts to external input. A systematic way to study the network response is to apply an oscillatory external input with fixed frequency *ω*, phase, and amplitude and observe the emerging frequency, phase, and amplitude of the output. If the amplitude of the external input is small compared to the stationary input, the network responds in a linear fashion: it only modifies phase and amplitude, while the output frequency equals the input frequency. This relationship is captured by the input-output transfer function *N* (*ω*) (Brunel et al., 2001; Brunel and Hakim, 1999; Lindner and Schimansky-Geier, 2001) which describes the frequency-dependent modulation of the output firing rate

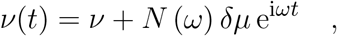

where *δμ* is the amplitude of the modulation to the input in units of Volt. The transfer function *N*(*ω*) is a complex function: Its absolute value describes the relative modulation of the firing rate. Its phase describes the phase shift that occurs between input and output. Since response properties are dominated by the modulation to the mean input, we here disregard a modulation of the noise intensity and denote the transfer function for a network of LIF neurons with instantaneous synapses as

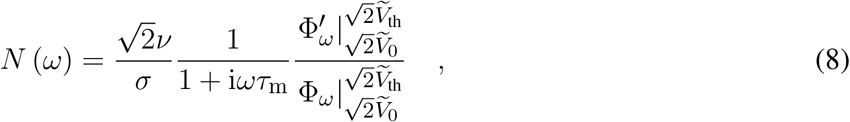

with the rescaled reset- and threshold-voltages 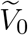 and 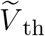 as defined in Eq. (3) and 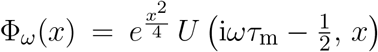 using the parabolic cylinder functions 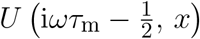 as defined in (Abramowitz and Stegun, 1974, Section 19.3) and (Olver et al., 2021, Section 12.2). 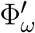 and 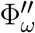 denote the first and second derivative by *x*, respectively. The transfer function given in Schuecker et al. (2015, Eq. 29) relates to our expression in Eq. (8) as follows: Our *N* (*ω*) *δμ* is equal to their *n_G_* (*ω*) *ν* implying that our *δμ* corresponds to their *ϵμ*; we assume their *n_H_* (*ω*) = 0. Besides, we swap the voltage boundaries both in numerator and denominator which cancels out.

For a neuronal network of LIF neurons with exponentially shaped post-synaptic currents, Schuecker et al. (2014, 2015) show that an analytical approximation of the colored-noise transfer function can be obtained by a shift of integration boundaries akin to Eq. (7):

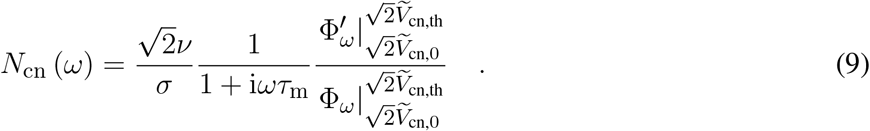

To take into account that, in neuronal networks, the current in Ampere is perturbed by the synaptic input, we include an additional low-pass filtering with the synaptic time constant into the definition:

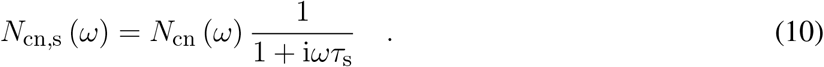

If the synaptic time constant is much smaller than the membrane time constant (*τ*_s_ ≪ *τ*_m_), an equivalent expression for the transfer function is obtained by a Taylor expansion around the original boundaries (cf. Schuecker et al. 2015, Eq. 30). The toolbox implements both variants and offers choosing between them by setting the argument `method` of `nnmt.lif.exp.transfer_function` to either `shift` or `taylor`.

Here, we demonstrate how to calculate the analytical “shift version” of the transfer function for different means and noise intensities of the input current (see **Figure 4**) and thereby reproduce **Figure 4** in Schuecker et al. (2015).

The crucial parts for producing **Figure 4** using NNMT are shown in **Listing 3** for one example combination of mean and noise intensity of the input current. Instead of using the model workflow with `nnmt.lif.exp.transfer_function`, we here employ the basic workflow, using `nnmt.lif.exp._transfer_function_shift` directly. This allows changing the mean input and its noise intensity independently of a network model’s structure, but requires two additional steps: First, the necessary parameters are loaded from a `yaml` file, converted to SI units and then stripped off the units using the utility function `nnmt.utils._convert_to_si_and_strip_units`. Second, the analysis frequencies are defined manually. In this example we choose logarithmically spaced frequencies, as we want to plot the results on a log-scale. Finally, the complex-valued transfer function is calculated and then split into its absolute value and phase. **Figure 4** shows that the transfer function acts as a low-pass filter that suppresses the amplitude of high frequency activity, introduces a phase lag, and can lead to resonance phenomena for certain configurations of mean input current and noise intensity.

The replication of the results from Schuecker et al. (2015) outlined here is also used in the integration tests of toolbox. Note that the implemented analytical form of the transfer function by Schuecker et al. (2015) is an approximation and deviations from a simulated ground truth are expected for higher frequencies (*ω*/2*π* ≳ 100 Hz at the given parameters).

**Listing 3:**
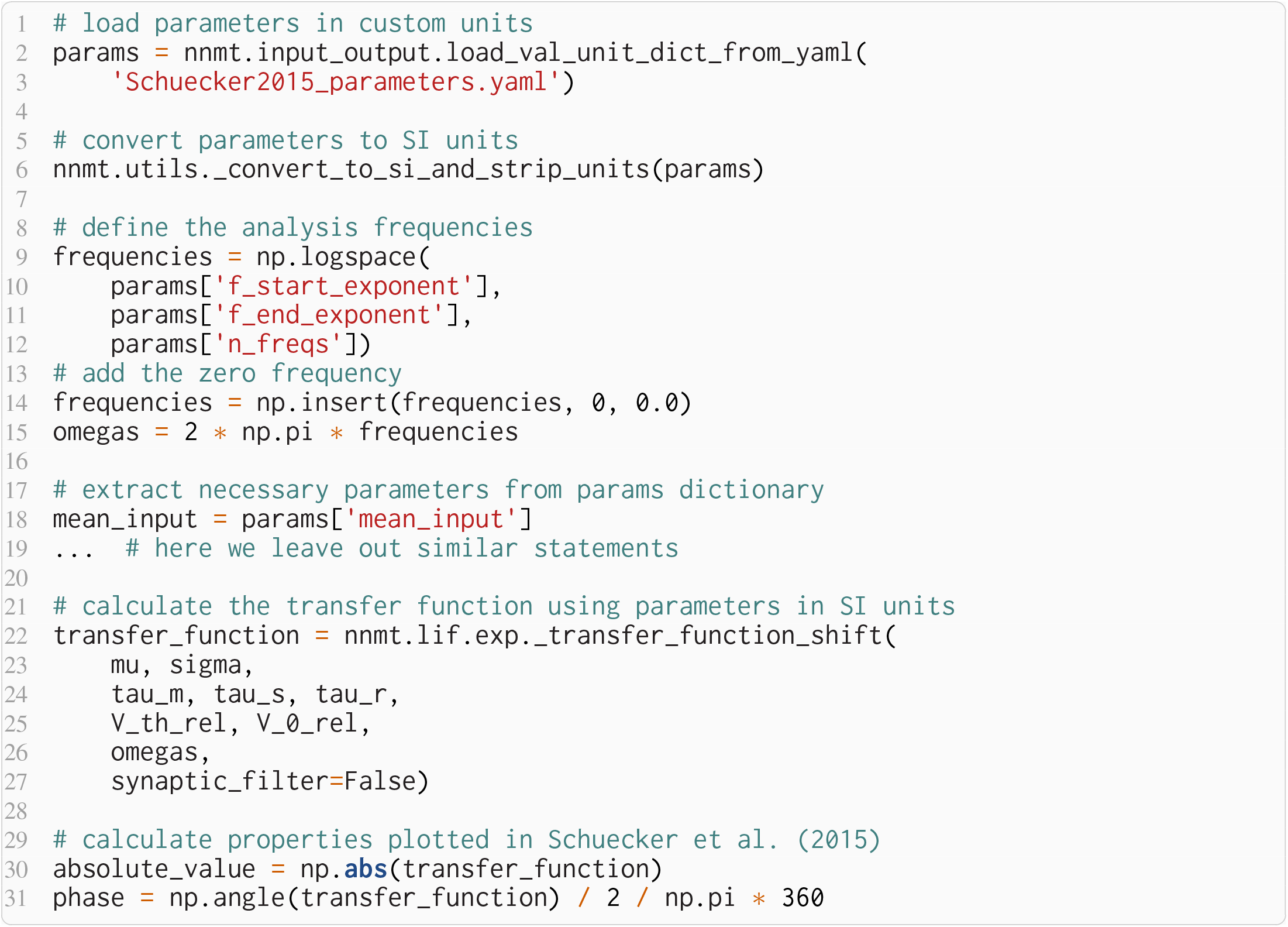
Example script for computing a transfer function shown in **Figure 4** using the method of shifted integration boundaries.

#### 3.3.2 Power Spectrum

Another dynamical property that is studied frequently is the power spectrum, which describes how the power of a signal is distributed across its different frequency components. This property for example reveals oscillations of the population activity. The power can be read off the diagonal entries of the crosscorrelations between population activities in the Fourier domain. The cross-correlations are determined by the network architecture, the stationary firing rates, and the network’s transfer function. The mean-field approximation of the power spectrum is obtained by studying the linear response of the network activity to the fluctuations caused by its spiking activity (Bos et al., 2016). The resulting analytical expression for the power spectra at a given angular frequency *ω* is

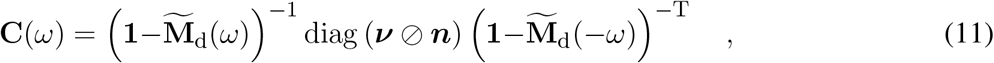

with ∅ denoting the elementwise (Hadamard) division, the effective connectivity matrix 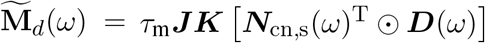, where ⊙ denotes the elementwise (Hadamard) product, the mean population firing rates ***ν***, and the numbers of neurons in each population ***n***. This effective connectivity combines the static, anatomical connectivity ***JK***, represented by synaptic weight matrix ***J*** and in-degree matrix ***K***, and dynamical quantities, represented by the transfer functions ***N***_cn,s_ (*ω*) (Eq. (10)) and the delay distributions ***D***(*ω*), both in Fourier domain.

The modular structure in combination with the model workflow of this toolbox permits a step-by-step calculation of the power spectra, as shown in **Listing 4**. The inherent structure of the theory is emphasized in these steps: After instantiating the network model class with given network parameters, the working point can be calculated, which includes calculating the population firing rates and the input statistics. This is necessary for determining the transfer functions. The calculation of the delay distribution matrix is then required for calculating the effective connectivity and to finally get an estimate of the power spectra. **Figure 5** reproduces **Figure 1E** in Bos et al. (2016) and shows the spectra for each population of the adjusted version (see **Table 1**) of the microcircuit model.

**Figure 5.**
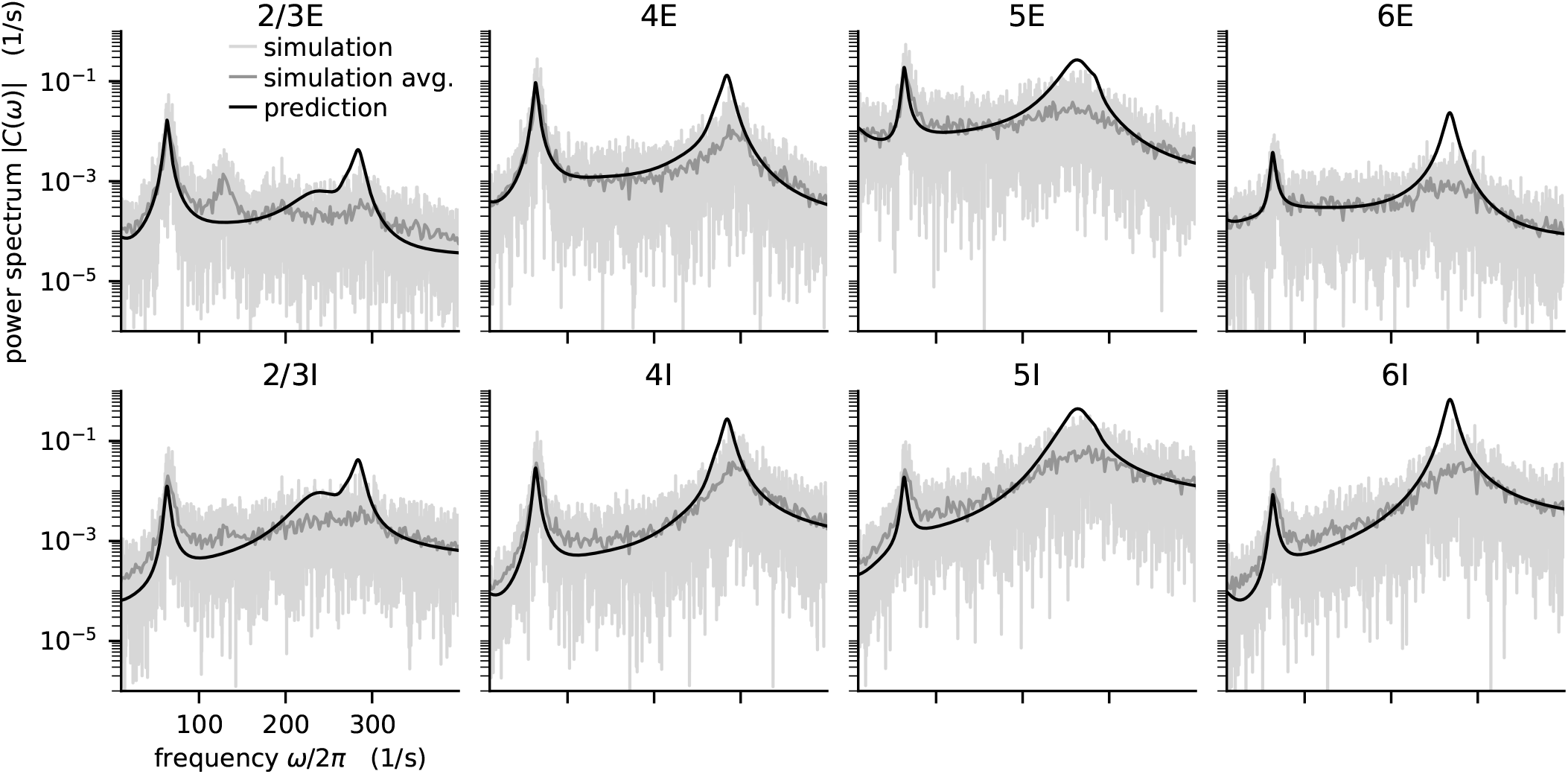
Power spectra of the population spiking activity in the adapted cortical microcircuit from Bos et al. (2016). The spiking activity of each population in a 10 s simulation of the model is binned with 1 ms resolution and the power spectrum of the resulting histogram is calculated by a fast Fourier transform (FFT; light gray curves). In addition, the simulation is split into 500 ms windows, the power spectrum calculated for each window and averaged across windows (gray curves). Black curves correspond to analytical prediction obtained with NNMT as described in **Listing 4**.

The numerical predictions obtained from the toolbox coincide with simulated data taken from the original publication (Bos et al., 2016) and reveal dominant oscillations of the population activities in the low-*γ* range around 64 Hz. Furthermore, faster oscillations with peak power around 300 Hz are predicted with higher magnitudes in the inhibitory populations 4I, 5I, and 6I.

#### 3.3.3 Sensitivity Measure

The power spectra shown in the previous section exhibit prominent peaks at certain frequencies, which indicate oscillatory activity. Naturally, this begs the question: which mechanism causes these oscillations? Bos et al. (2016) expose the crucial role that the microcircuit’s connectivity plays in shaping the power spectra of this network model. They have developed a method called *sensitivity measure* to directly relate the influence of the anatomical connections, especially the in-degree matrix, on the power spectra.

The power spectrum of the *i*-th population *C_ii_*(*ω*) receives a contribution from each eigenvalue *λ_j_* of the effective connectivity matrix, *C_ii_*(*ω*) ∝ 1/(1 – λ_*j*_ (*ω*))^2^. Such a contribution consequently diverges as the complex-valued λ_*j*_ approaches 1 + 0i in the complex plane, which is referred to as the point of instability. This relation can be derived by replacing the effective connectivity matrix 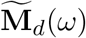 in Eq. (11) by its eigendecomposition. The sensitivity measure leverages this relationship and evaluates how a change in the in-degree matrix affects the eigenvalues of the effective connectivity and thus indirectly the power spectrum. Bos et al. (2016) introduce a small perturbation *α_kl_* of the in-degree matrix, which allows writing the effective connectivity matrix as 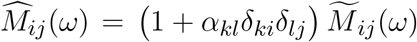, where we dropped the delay subscript *d*. The sensitivity measure *Z_j,kl_* (*ω*) describes how the *j*-th eigenvalue of the effective connectivity matrix varies when the *kl*-th element of the in-degree matrix is changed

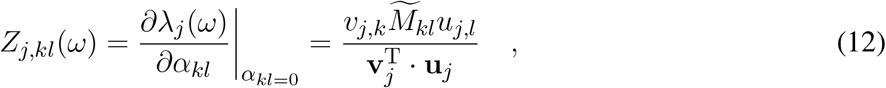

where 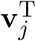 and **u**_*j*_ are the left and right eigenvectors of 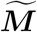 corresponding to eigenvalue λ_*j*_ (*ω*).

**Listing 4:**
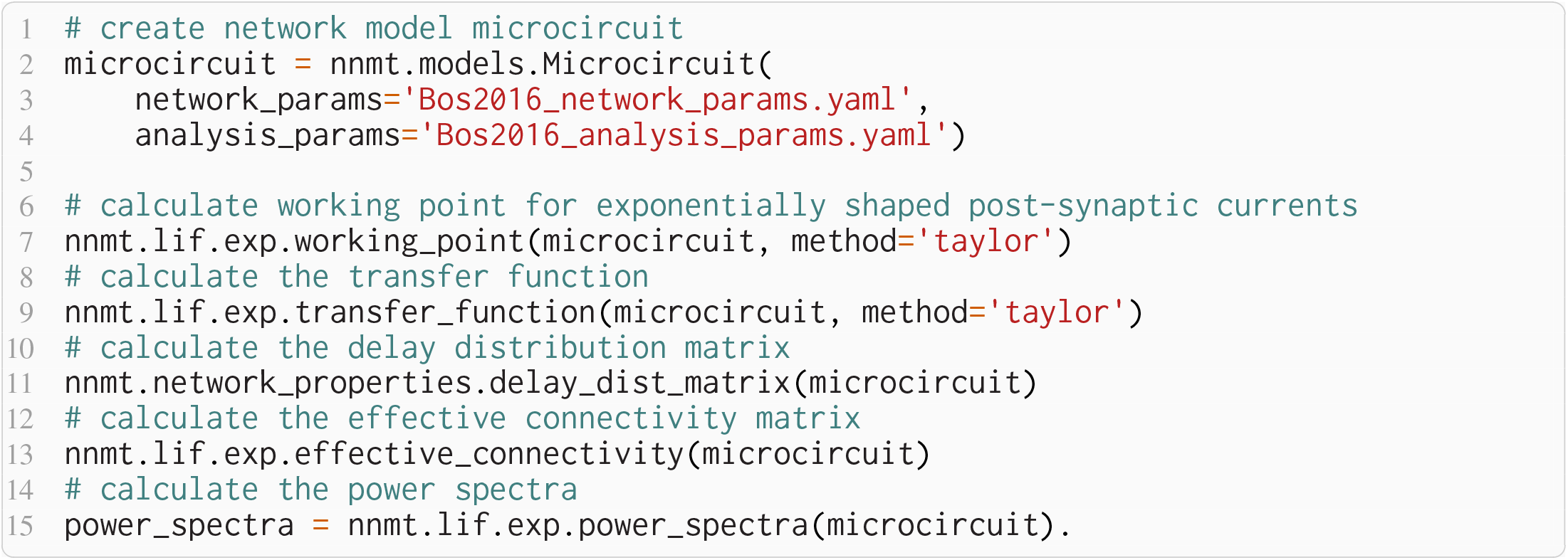
Example script to produce the data shown in **Figure 5**.

The complex sensitivity measure can be understood in terms of two components: 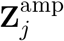 is the projection of the matrix **Z**_*j*_ onto the direction in the complex plane defined by 1 – λ_*j*_ (*ω*); it describes how, when the in-degree matrix is perturbed, the complex-valued λ_*j*_ (*ω*) moves towards or away from the instability 1 + 0*i*, and consequently how the amplitude of the power spectrum at frequency *ω* increases or decreases. 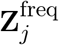 is the projection onto the perpendicular direction and thus describes how the peak frequency of the power spectrum changes with the perturbation of the in-degree matrix.

The toolbox makes this intricate measure accessible by supplying two tools: After computing the required working point, transfer function, and delay distribution, the tool `nnmt.lif.exp.sensitivity_measure` computes the sensitivity measure at a given frequency for one specific eigenvalue. By default, this is the eigenvalue which is closest to the instability 1 + 0*i*. To perform the computation, we just need to add one line to **Listing 4**:

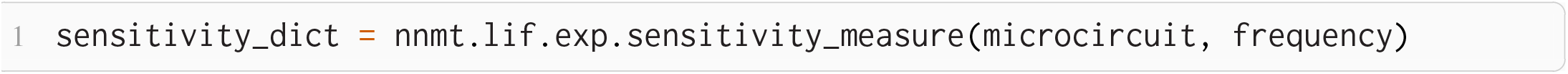

The result is returned in form of a dictionary that contains the sensitivity measure and its projections. The tool `nnmt.lif.exp.sensitivity_measure_per_eigenmode` wraps that basic function and calculates the sensitivity measure for all eigenvalues at the frequency for which each eigenvalue is closest to instability.

According to the original publication (Bos et al., 2016), the peak around 300 Hz has contributions from four different eigenvalues of the effective connectivity matrix. **Figure 6** shows the projections of the sensitivity measure at the frequency for which the corresponding eigenvalues are closest to the instability, as in Figure 4 of Bos et al. (2016). The sensitivity measure returns one value for each connection between populations in the network model. For 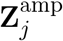 a negative value indicates that increasing the in-degrees of a specific connection causes the amplitude of the power spectrum at the evaluated frequency to drop. If the value is positive, the amplitude is predicted to grow as the in-degrees increase. Similarly, for positive 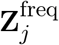 the frequency of the peak in the power spectrum shifts towards higher values as in-degrees increase, and vice versa. The main finding in this analysis is that the high-*γ* peak seems to be affected by inhibitory self-connections of each population.

**Figure 6.**
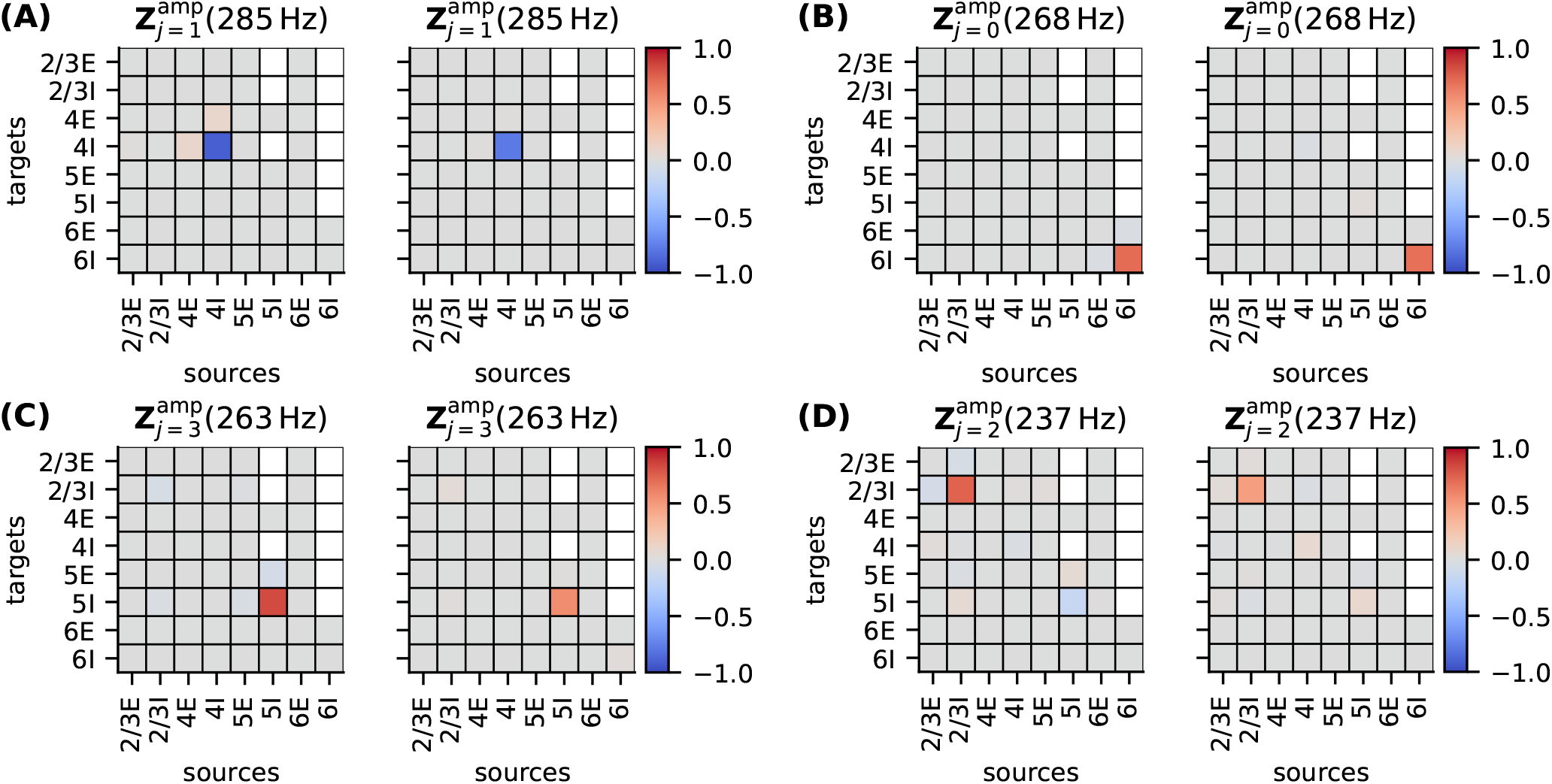
Sensitivity measure of four eigenmodes of the effective connectivity relevant for high-*γ* oscillations. The sensitivity measure for each mode is evaluated at the frequency where the corresponding eigenvalue is closest to the point of instability 1 + 0*i* in complex plane. 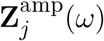 visualizes the influence of a perturbation of a connection on the peak amplitude of the power spectrum. 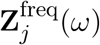 shows the impact on the peak frequency. Non-existent connections are masked white.

To decrease the high-*γ* peak in the power spectrum, one could therefore increase the 4I to 4I connections (cp. panel **(A)** in **Figure 6**)

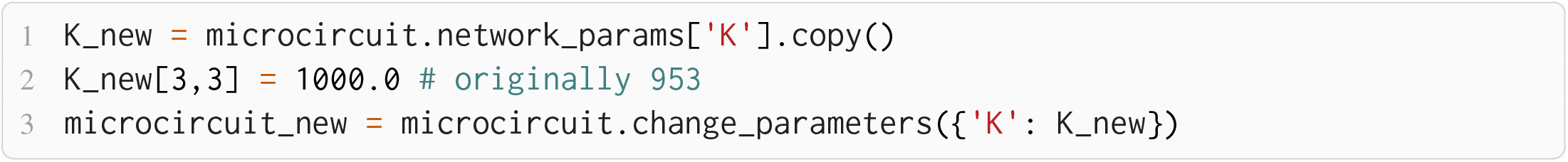

and calculate the power spectrum as in **Listing 4** again to validate the change.

### 3.4 Fitting Spiking to Rate Model and Predicting Pattern Formation

If the neurons of a network are spatially organized and connected according to a distance-dependent profile, the spiking activity may exhibit pattern formation in space and time, including wave-like phenomena. Senk et al. (2020) set out to scrutinize the non-trivial relationship between the parameters of such a network model and the emerging activity patterns. The model they use is a two-population network of excitatory E and inhibitory I spiking neurons, illustrated in **Figure 7**. All neurons are of type LIF with exponentially shaped post-synaptic currents. The neuron populations are recurrently connected to each other and themselves and they receive additional external excitatory E_ext_ and inhibitory I_ext_ Poisson spike input of adjustable rate as shown in **Figure 7A**. The spatial arrangement of neurons on a ring is illustrated in **Figure 7B** and the boxcar-shaped connectivity profiles in **Figure 7C**.

**Figure 7.**
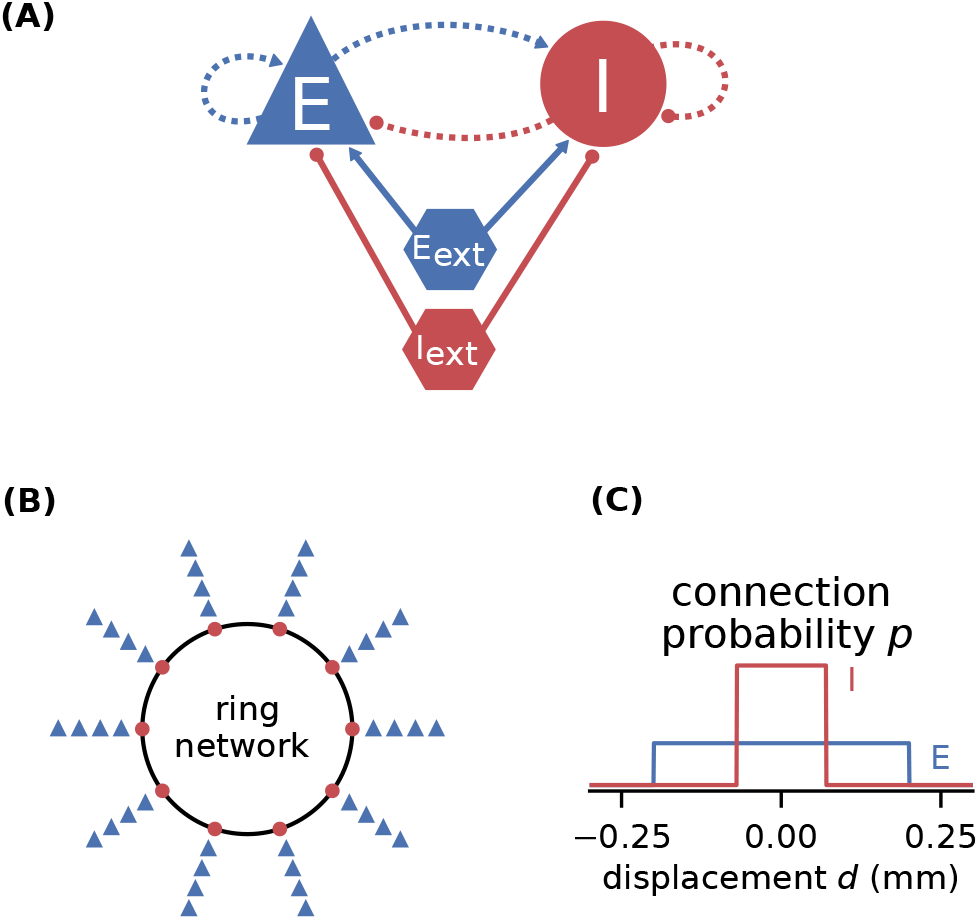
Illustrations of spiking network model by Senk et al. (2020). **(A)** Excitatory and inhibitory neuronal populations randomly connected with fixed in-degree and multapses allowed (autapses prohibited). External excitatory and inhibitory Poisson drive to all neurons. Same notation as in **Figure 2A**. **(B)** One inhibitory and three excitatory neurons per grid position on a one-dimensional domain with periodic boundary conditions (ring network). **(C)** Normalized, boxcar-shaped connection probability with wider excitation than inhibition. For model details and parameters, see Tables II–IV of Senk et al. (2020); the specific values given in the caption of their Figure 6 are used throughout here.

In the following, we consider a mean-field approximation of the spiking model with spatial averaging, that is a time and space continuous approximation of the discrete model as derived in Senk et al. (2020, E. Linearization of spiking network model). We demonstrate three methods used in the original study: First, Section 3.4.1 explains how a model can be brought to a defined state characterized by its working point. The working point is given by the mean *μ* and noise intensity *σ* of the input to a neuron, which are both quantities derived from network parameters and require the calculation of the firing rates. With the spiking model in that defined state, Section 3.4.2 then maps its transfer function to the one of a rate model. Section 3.4.3 finally shows that this working-point dependent rate model allows for an analytical linear stability analysis of the network accounting for its spatial structure. This analysis can reveal transitions to spatial and temporal oscillatory states which, when mapped back to the parameters of the spiking model, may manifest in distinct patterns of simulated spiking activity.

#### 3.4.1 Setting the working point by changing network parameters

With network and analysis parameters predefined in `yaml` files, we set up a network model using the example model class `Basic`:

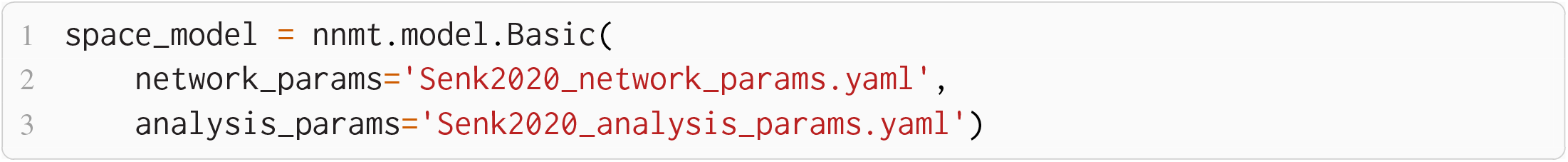

Upon initialization the given parameters are automatically converted into the format used by NNMT’s tools. For instance, relative spike reset and threshold potentials are derived from the absolute values, connection strengths in units of volt are computed from the post-synaptic current amplitudes in ampere, and all values are scaled to SI units.

We aim to bring the network to a defined state by fixing the working point but also want to explore if the procedure of fitting the transfer function still works for different network states. For a parameter space exploration, we use a method to change parameters provided by the model class and scan through a number of different working points of the network. To obtain the required input for a target working point, we adjust the external excitatory and inhibitory firing rates accordingly; NNMT uses a vectorized version of the equations given in Senk et al. (2020, Appendix F) to calculate the external rates needed:

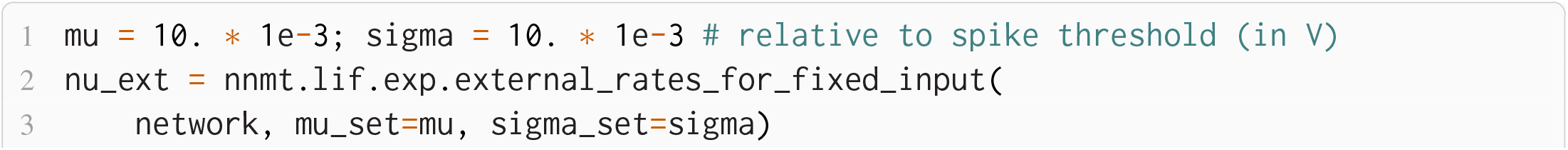
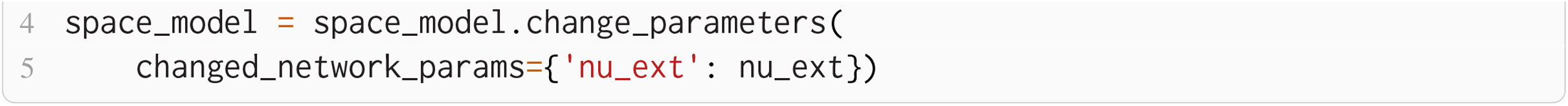

The resulting external rates for different choices of (*μ*, *σ*) are color-coded in the first two plots of **Figure 8A**. The third plot shows the corresponding firing rates of the neurons, which are stored in the results of the model instance when computing the working point explicitly:

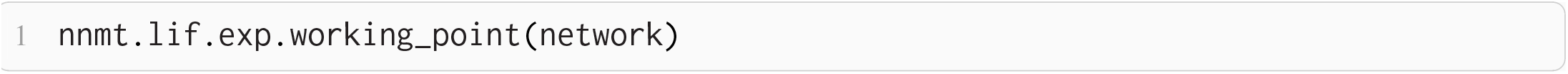

**Figure 8.**
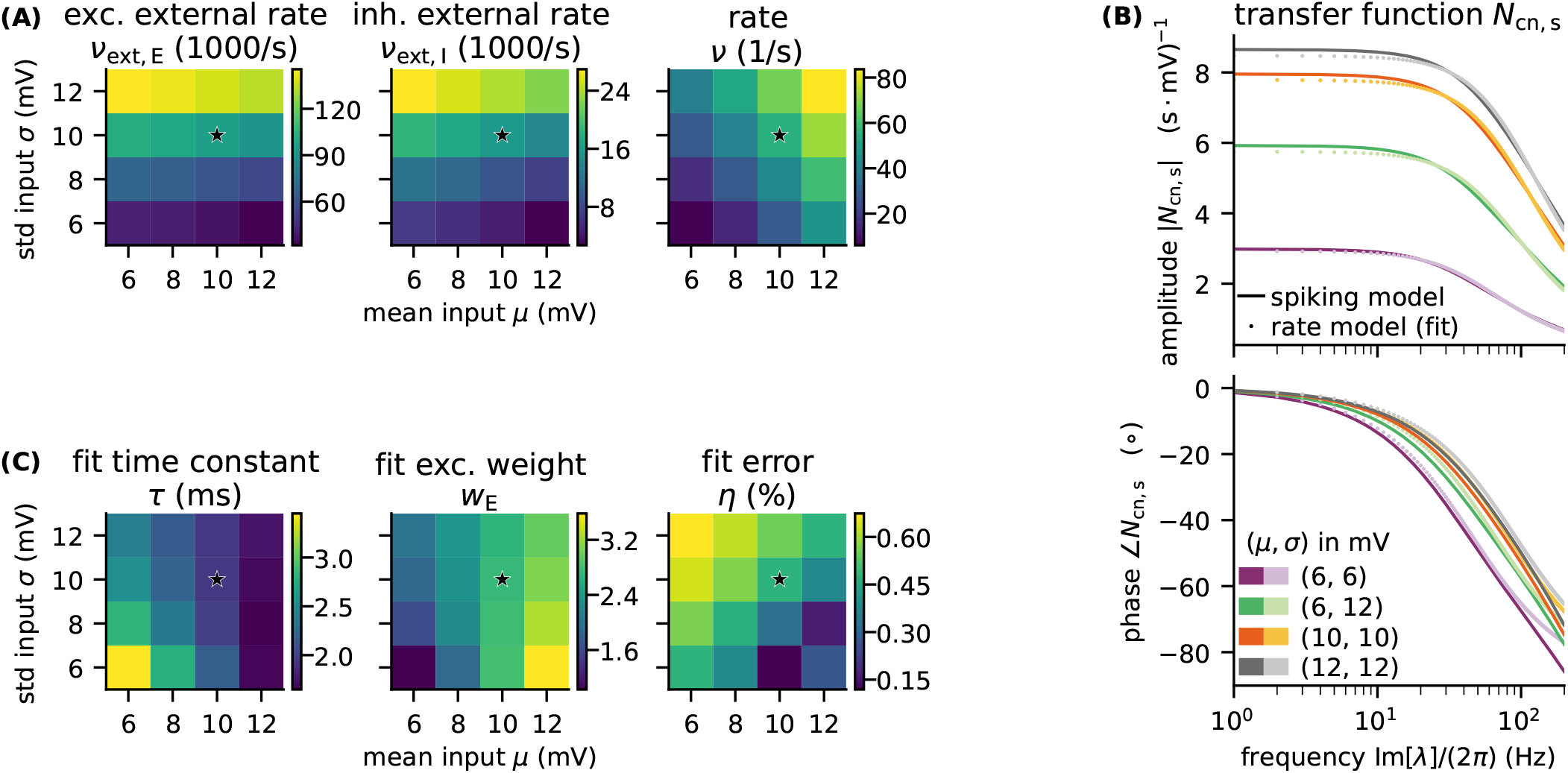
Network parameters and mean-field results from scanning through different working points. Working point (*μ*, *σ*) combines mean input *μ* and noise intensity of input *σ*. **(A)** External excitatory *ν*_ext,E_ and inhibitory *ν*_ext,I_ Poisson rates required to set (*μ*, *σ*) and resulting firing rates *ν*. **(B)** Transfer function *N*_cn,s_ of spiking model and fitted rate-model approximation with low-pass filter for selected (*μ*, *σ*) (top: amplitude, bottom: phase). **(C)** Fit results (time constants *τ* and excitatory weights *w*_E_) and fit errors *η*. The inhibitory weights are *w*_I_ = –*gw*_E_ with *g* = 5. Star marker in panels (A) and (C) denotes target working point (10, 10) mV. Similar displays as in Senk et al. (2020, Figure 5).

#### 3.4.2 Parameter mapping by fitting the transfer function

We map the parameters of the spiking model to a corresponding rate model by, first, computing the transfer function *N*_cn,s_ given in Eq. (10) of the spiking model, and second, performing a least-squares fit to the simpler transfer function of the rate model, for details see Senk et al. (2020, F Comparison of neural-field and spiking models). The dynamics of the rate model can be written as a convolution equation with the temporal kernel Θ (*t*) · *w*/*τ* · exp (–*t*/*τ*) using the Heaviside function Θ; *τ* is the time constant and w the unitless weight. Hence, the transfer function is just the one of a low-pass filter, *N*_LP_ = *w*/(1 + λ*τ*), where λ is the frequency in Laplace domain. The tool to fit the transfer function requires that the actual transfer function has been computed beforehand and returns the fit for the same frequencies together with *τ*, *w*, and the combined fit error *η*:

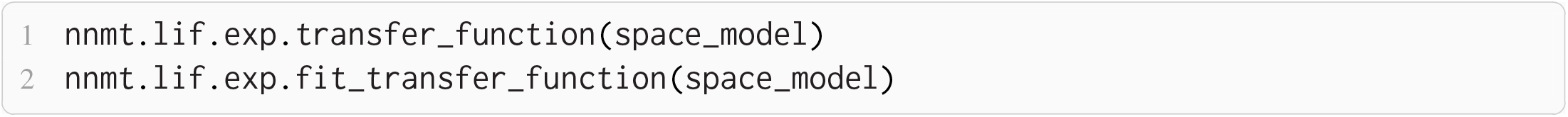

**Figure 8B** illustrates the amplitude and phase of the transfer function and its fit for a few (*μ*, *σ*) combinations. The plots of **Figure 8C** show the fitted time constants, the fitted excitatory weight, and the combined fit error. The inhibitory weight is proportional to the excitatory one in the same way as the post-synaptic current amplitudes.

#### 3.4.3 Linear stability analysis of spatially structured model with delay

The rate model with fitted parameters combined with the spatial connectivity profile is considered a neural field model, which lends itself to analytical linear stability analysis, as described in detail in Senk et al. (2020, A. Linear stability analysis of a neural-field model). In brief, this analysis seeks for pattern-forming instabilities or, in other words, for transitions from homogeneous steady states to oscillatory states as a function of a bifurcation parameter, here the delay *d*. The complex-valued, temporal eigenvalue λ of the linearized time-delay system is an indicator for the system’s overall stability and can serve as a predictor for temporal oscillations, whereas the spatial oscillations are characterized by the real-valued wave number *k*. Solutions that relate λ and *k* with the model parameters are given by a characteristic equation, which in our case reads (Senk et al., 2020, Eq. 7):

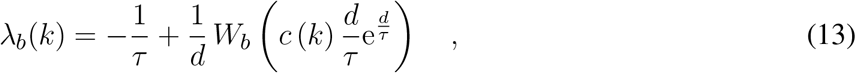

with the time constant of the rate model **τ**, the multi-valued Lambert *W_b_* function on branch *b* (Corless et al., 1996), and the effective connectivity profile *c*(*k*), which combines the weights *w* and the Fourier transforms of the boxcar-shaped spatial connectivity profiles. NNMT provides an implementation to solve this characteristic equation in its `linear_stability` module using the `spatial` module for the profile:

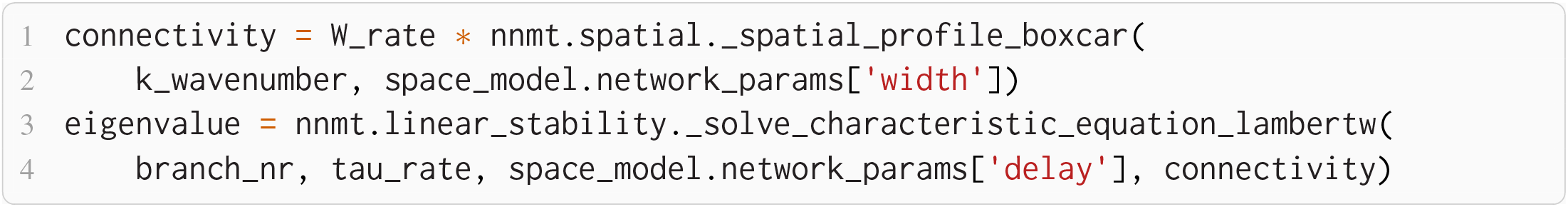

**Figure 9A** shows that the computed eigenvalues come for the given network parameters in complex conjugate pairs. Temporal oscillations are expected to occur if the real part of the eigenvalue on the largest branch becomes positive; the oscillation frequency can then be read off the imaginary part of that eigenvalue. In this example, the largest eigenvalue λ* on the principle branch (*b* = 0) has a real part that is just above zero. The respective wave number *k*\* is positive, which indicates spatial oscillations as well. The oscillations in both time and space predicted for the rate model imply that the activity of the corresponding spiking model might exhibit wave trains, i.e., temporally and spatially periodic patterns. The predicted propagation speed of the wave trains is given by the phase velocity Im [λ*] /*k*\*.

**Figure 9.**
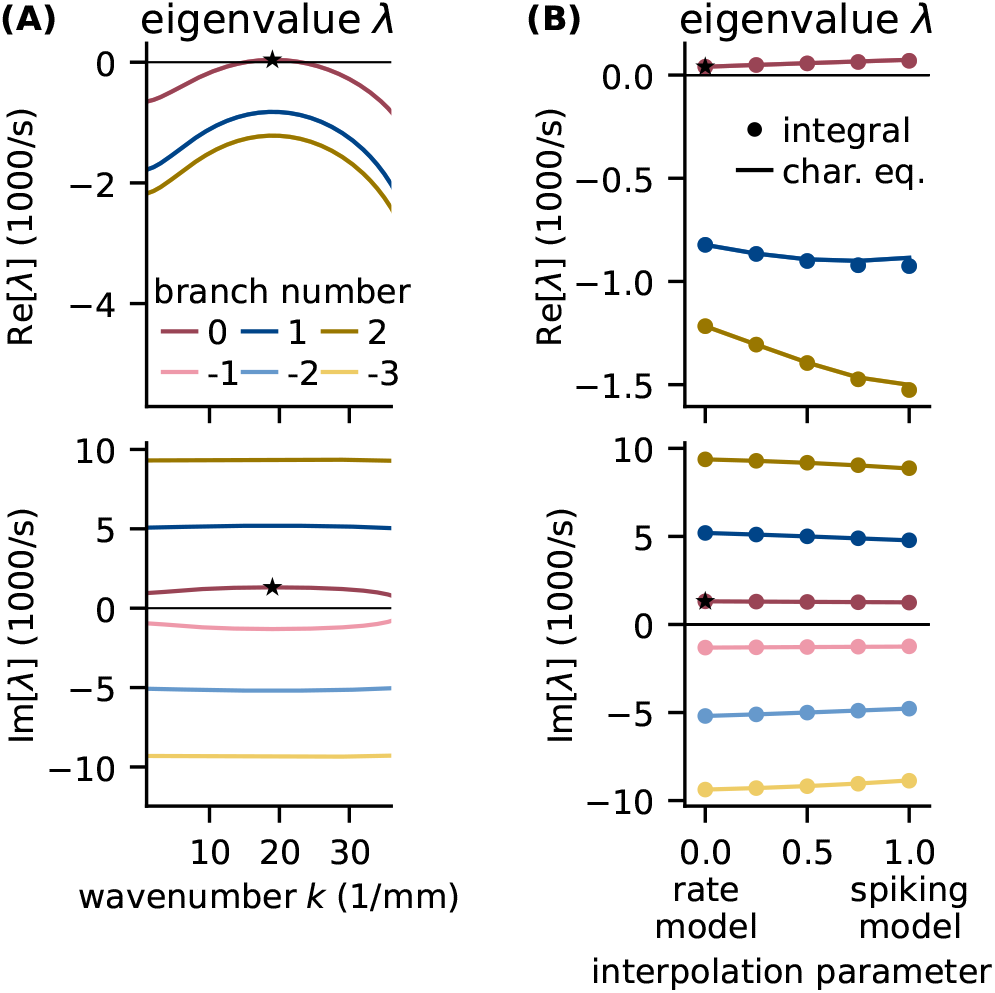
Linear stability analysis of spatially structured network model. **(A)** Analytically exact solution for real (top) and imaginary (bottom) part of eigenvalue λ versus wavenumber *k* using rate model derived by fit of spiking model at working point (*μ*, *σ*) = (10, 10) mV. Color-coded branches of Lambert W function; maximum real eigenvalue (star marker) on principal branch (*b* = 0). **(B)** Linear interpolation between rate (*α* = 0) and spiking model (*α* = 1) by numerical integration of Senk et al. (2020, Eq. 30) (solid line) and by numerically solving the characteristic equation in Senk et al. (2020, Eq. 29) (circular markers). Star markers at same data points as in panel (A). Similar displays as in Senk et al. (2020, Figure 6).

To determine whether the results obtained with the rate model are transferable to the spiking model, **Figure 9B** interpolates the analytical solutions of the rate model (*α* = 0) to solutions of the spiking model (*α* = 1), which can only be computed numerically; for details see Senk et al. (2020, F.2 Linear interpolation between the transfer functions). Since the eigenvalues estimated this way show only little differences between rate and spiking model, we conclude that predictions from the rate model should hold also for the spiking model in the parameter regime tested. Following the argument of Senk et al. (2020), the predicted pattern formation could next be tested in a numerical simulation of the discrete spiking network model (for such results with the parameters used here, see their Figure 10(c) for *d* = 1.5 ms).

## 4 DISCUSSION

Mean-field theory grants important insights into the dynamics of neuronal networks. However, the lack of a publicly available numerical implementation for most methods entails a significant initial investment of time and effort prior to any scientific investigations, thereby slowing down progress.

This paper describes the open-source toolbox NNMT that provides access to various neuronal network model analyses based on mean-field theory. As use cases, we reproduce known results from the literature: The non-linear relation between the firing rates and the external input of an E-I-network (Sanzeni et al., 2020). The firing rates of a cortical microcircuit model, its response to oscillatory input, its power spectrum, and the identification of the connections that predominantly contribute to the model’s high frequency oscillations (Bos et al., 2016; Schuecker et al., 2015). Finally, pattern formation in a spiking network, analyzed by mapping it to a rate model and conducting a linear stability analysis (Senk et al., 2020).

Here, we start with a general discussion on the benefits and drawbacks of collecting methods in a common toolbox. We expand on further use cases for NNMT and also point out the inherent limitations of analytical methods for neuronal network analysis. We conclude with suggestions on how NNMT might overcome some of these limitations and how the toolbox may grow and develop in the future.

### 4.1 Using a Toolbox vs. Writing Your Own Code

Why is there not yet a toolbox like this (of which we are aware of)? And consequently, is there actually a need for a toolbox like this? Here, we would like to share our thoughts on these questions (see also Riquelme and Gjorgjieva, 2021).

Without a toolbox, everybody who wants to use an analytical method for network model analysis, needs to implement them on their own. This has an advantage: a detailed understanding of the theory behind the method and the implementation itself. Accordingly, a scientist following this path will know exactly how to apply the method and, most importantly, when the method cannot be applied and conversely when it will lead to misleading or wrong results. Providing those tools in a ready-to-use fashion comes with the danger of enticing users into applying those tools without understanding their limitations. And this might lead them to draw false conclusions based on the results they get from using the toolbox. This can only be avoided by making the users aware of the limitations a method comes with and providing sufficient information in an easy-to-find and comprehensible form. Therefore, in NNMT, we try to address this issue by raising warnings whenever the valid parameter regimes are left and referring the user to the package’s documentation, where we try our best to make all the necessary information available. Of course, the best protection against running into those problems in the first place is reading the documentation before using the tools.

However, implementing everything on your own also has disadvantages: It takes time and therefore money, which possibly is invested into the same tool over and over again by different groups around the world; which also can be seen as a waste of resources. Often, implementing and debugging analytical tools is not straightforward and can be a matter of several weeks. This naturally implicates a certain complexity barrier, beyond which such tools, which might be dependent on other tools, cannot develop, as the effort of implementing them becomes too large. In a similar vein, which kind of questions researchers pursue is limited by and therefore depends on the tools they have at hand. The availability of sophisticated neural network simulators for example has lead to the development of conceptually new and more complex neural network models, precisely because their users did not have to bother about implementation details any longer and could instead focus on actual research questions. We hope that collecting useful implementations of analytical tools for network model analysis will have a similar effect on the development of new tools and that it might lead to new, creative ways of applying them.

Finally, having an open-source toolbox, everybody can contribute to, has the compelling advantage of leveraging the knowledge of a large part of the scientific community. Such a toolbox can serve as a platform for collecting standard implementations of methods, which have been tested thoroughly and have been validated by many users. Of course, if a toolbox does not have the functionality a user might need, it needs some extra effort to first learn how the toolbox works and to find out where necessary changes need to be made. But instead of coding everything from scratch, it might only be a few lines that need to be modified. And most importantly, in the end not only the own lab but the whole community may benefit from this extra effort.

### 4.2 Use Cases

In Section 3, we present concrete examples of how to apply some of the tools available. Here, we discuss further, more general use cases for NNMT: parameter space exploration and fast prototyping, developing an intuitive understanding of network models, extracting network mechanisms, investigating hypotheses using limited computational resources, and hybrid approaches.

Analytical methods have the advantage of being fast, and typically they only require a limited amount of computational resources. All tools currently implemented in NNMT can be run on a notebook without any problems. And the computational costs for calculating analytical estimates of dynamical network properties like firing rates, as opposed to the costs of running simulations of a network model, are independent of the number of neurons the network is composed of. This becomes especially relevant if one wants to do parameter space explorations, for which many simulations have to be performed. To speed up prototyping, a modeler can first perform a parameter scan using analytical tools from NNMT to get an estimate of the right parameter regimes and subsequently run simulations on this restricted set of parameters to arrive at the final model parameters.

But additionally to speeding up undirected parameter space explorations, analytical methods might guide parameter space explorations in another way: namely, by giving some intuitive understanding of the relation between network model parameters and network dynamics. The prime example implemented in NNMT is the sensitivity measure presented in Section 3.3.3, which provides a “frequency-dependent connectivity map that reveals connections crucial for the peak amplitude and frequency of the observed oscillations [of a network model] and identifies the minimal circuit generating a given frequency” (Bos et al., 2016). This illustrates a case in which a mean-field method can inform the modeler about the origin of a model’s dynamics, and it identifies which model parameters need to be adjusted to modify that dynamics.

Similarly, a modeler might be interested in extracting network mechanisms, answering the question, which features of a network cause some dynamics to happen. We have included a method for mapping the parameters of a spiking network model to a rate network model, which is presented in Section 3.4. Using this method, one can check whether the dynamical properties under investigation are still present in a simulation of a simpler rate network. Consequently, one can conclude whether spiking dynamics is a crucial component for the occurrence of the respective dynamical property, or whether it is sufficient to study the simpler model. This way, NNMT can help identifying the essential components that lead to some dynamical features.

Another way to take advantage of analytical tools, is to investigate hypotheses directly, employing those methods. In Section 3.2.1 the toolbox is used to show non-trivial effects regarding the firing rates. Users could first perform a similar analysis for their research question, using the computationally cheap mean-field methods, thereby deepening their understanding of the problem, and then apply for compute time on a high performance cluster afterwards to check those predictions in simulations. Thus, NNMT can serve as an additional instrument for investigating research questions, especially for researchers who do not have free access to a high performance computer. Of course, the limitations explained in Section 4.3 always have to be taken into account to avoid misleading conclusions.

Finally, NNMT can be used for hybrid approaches, where one quantity is extracted from simulations or experiments, and the toolbox is then used to calculate another quantity that might not be directly accessible otherwise. Getting access to possibly abstract, but informative measures opens up the possibility to conduct new kinds of experiments and simulations, which investigate the behavior of such measures under possibly complex interventions that could not be done using analytical tools alone.

### 4.3 Limitations

NNMT has two major limitations: restrictions of valid parameter regimes due to necessary approximations in the analytics, and the restriction to network, neuron, and synapse models, as well as observables, for which a mean-field theory exists.

Analytical methods can provide good estimates of properties of a network model. But it is important to state that those methods almost always depend on some kinds of approximations, which can only be justified if the network under consideration fulfills certain assumptions. If those assumptions are not met, at least approximately, one cannot assure that the corresponding analytical methods give valid results. For example, NNMT calculates the firing rates for leaky integrate-and-fire neurons with exponential synapses using the results from Fourcaud and Brunel (2002). This requires the neuronal populations to be large enough, it requires them to receive a high enough number of uncorrelated inputs, such that the central limit theorem can be applied, and the solution depends on the assumption that the synaptic time constant is small compared to the membrane time constant. Fortunately, those assumptions are known and some of them can be checked for. However, there are assumptions that cannot be tested, as they would necessitate a theory going beyond the mean-field approximation, such as the network state being sufficiently uncorrelated. If a tool from NNMT is applied outside of its valid parameter ranges, NNMT will raise a warning, informing the user about the issue, whenever this is possible.

To better understand the sources of inaccuracies in NNMT, it is helpful to consider a comparison to direct simulations of network models. Simulations only suffer from inaccuracies due to the numerical implementation, which can be improved upon by using enhanced numerical schemes or simply choosing smaller integration steps. Numerical inaccuracies occur in NNMT as well, and they can be remedied in a similar fashion. We address this problem with an extensive test suite based on previously published results. But often, the assumptions discussed in the previous paragraph cannot be met perfectly. For example some analytical estimates might only be exact if the number of neurons in the network is infinite. Obviously this cannot be achieved in a realistic network. Therefore, the approximations applied in analytical methods can introduce a second source of inaccuracy, which originates from neglecting higher order contributions. This kind of inaccuracy cannot be resolved by improving some hyperparameters, as it depends directly on the choice of network parameters. This explains the discrepancy between networks model simulations and analytical results. Nevertheless, analytical methods will lead to reliable estimates if applied within the valid parameter regimes (for example see **Figure 3B**).

Finally, mean-field methods will always be restricted to network, neuron, and synapse models for which the relation between model parameters and activity statistics can be derived. If the model under consideration does not allow deriving such a relation, resorting to a similar model, as done in Section 3.2.2, may be a viable solution. But if a model becomes too complex, it is usually impossible to derive any analytical results. In such cases, one might prefer simulations (Einevoll et al., 2019). However, analytical methods come with advantages that no other approach can offer, as we have shown in Section 4.2. In the following section we explain how interested colleagues can support the community in fully utilizing these advantages.

### 4.4 How to Contribute and Outlook

As discussed in Section 2, NNMT has been designed for modularity, flexibility, and extensibility. The aim is to provide a platform which can fit many different mean-field methods. NNMT is an open-source toolbox, and we would like to encourage scientists to contribute by improving existing methods and sharing their own methods and implementations via the toolbox. This can be done via the standard pull request workflow on GitHub. If you are interested and would like to know more about the procedure, we refer you to the “Contributors guide” of the official documentation of NNMT^4^.

A toolbox like NNMT always is an ongoing project, and there are various aspects that can be improved. Here we shortly discuss three of them: parallelization, extension of valid parameter regimes, and different neuron types and network architectures.

NNMT in its current state is partly vectorized but it currently does not include parallelized methods, e.g., using multiprocessing or MPI for Python (`mpi4py`). Vectorization relies on `NumPy` (Harris et al., 2020) and `SciPy` (Virtanen et al., 2020) which are thread-parallel for specific backends, e.g., IntelMKL. With the tools available in the toolbox at the moment, run-time only becomes an issue in extensive parameter scans, for instance, when the transfer function needs to be calculated for a large range of frequencies. To further reduce the runtime, the code could be made fully vectorized. Alternatively, parallelization of many tools in NNMT is straightforward, as many of them include `for` loops over the different populations of a network model and `for` loops over the different analysis frequencies.

As discussed in Section 4.3, many mean-field methods only give valid results for certain parameter ranges. But often, there exist different approximations for the same quantity, valid in complementary parameter regimes. A concrete example is the currently implemented version of the transfer function for leaky integrate-and-fire neurons, based on the work of Schuecker et al. (2015), which gives a good estimate for small synaptic time constants compared to the membrane time constant *τ*_s_/*τ*_m_ ≪ 1. A complementary estimate for long synaptic time constants *τ*_m_/*τ*_s_ ≪ 1, which could be implemented, has been developed by Moreno-Bote and Parga (2006). Similarly, the current implementation of the firing rates of leaky integrate-and-fire neurons is based on the work of Fourcaud and Brunel (2002), and only valid for *τ*_s_/*τ*_m_ ≪ 1. But van Vreeswijk and Farkhooi (2019) have developed a method accurate for all combinations of synaptic and membrane time constants. Adding such complementary and improved tools will enhance the applicability of the toolbox.

Finally, due to the research focus at our lab, NNMT presently mainly contains tools for LIF neurons. But the structure of NNMT allows for adding methods for different neuron types, like binary (Ginzburg and Sompolinsky, 1994) or conductance-based neurons (Izhikevich, 2007; Richardson, 2007). In a similar manner, most methods focus on networks with random connectivity, but we already added some tools for networks with spatially organized connectivity. We hope that in the future many scientists will contribute to this collection of analytical methods for neuronal network model analysis, such that at some point, we will have tools from all parts of mean-field theory of neuronal networks, made accessible in a usable format to all neuroscientists.

## AUTHOR CONTRIBUTIONS

HB and MH developed and implemented the code base and the inital version of the toolbox. ML, JS, and SE designed the current version of the toolbox. ML implemented the current version of the toolbox, vectorized and generalized tools, developed and implemented the test suite, wrote the documentation, and created the example shown in Section 3.2.2. AvM improved the numerics of the firing rate integration (Appendix Section 5.1) and created the example shown in Section 3.2.1. SE implemented integration tests, improved the functions related to the `sensitivity_measure`, and created the examples shown in Section 3.3. JS developed and implemented the tools used in Section 3.4 and created the respective example. ML, JS, SE, AvM, and MH wrote this article. All authors approved the submitted version.

## ACKNOWLEDGMENTS

We would like to thank Jannis Schuecker, who has contributed to the development and implementation of the code base and the inital version of the toolbox. This project has received funding from the European Union’s Horizon 2020 Framework Programme for Research and Innovation under Specific Grant Agreement No. 720270 (HBP SGA1), 785907 (HBP SGA2), and 945539 (HBP SGA3), and has been partially funded by the Deutsche Forschungsgemeinschaft (DFG, German Research Foundation) - 368482240/GRK2416. This research was supported by the Joint Lab “Supercomputing and Modeling for the Human Brain”.

## 5 APPENDIX

### 5.1 Siegert Implementation

Here, we describe how we solve the integral in Eq. (2) numerically in a fully vectorized manner. The difficulty in Eq. (2), 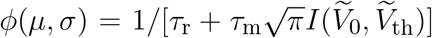 where 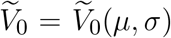 and 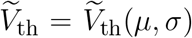 are determined by either Eq. (3) or Eq. (7), is posed by the integral

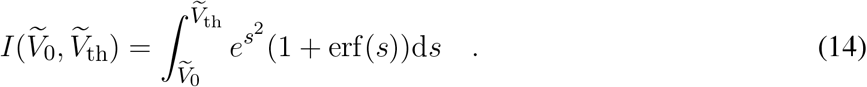

This integral is problematic due to the multiplication of 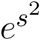 and 1 + erf (*s*) in the integrand which leads to overflow and loss of significance.

To address this, we split the integral into different domains depending on the sign of the integration variable. Furthermore, we use the scaled complementary error function

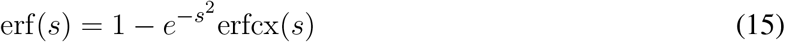

to extract the leading exponential contribution. Importantly, erfcx(*s*) decreases monotonically from erfcx(0) = 1 with power law asymptotics 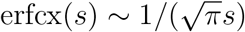, hence it does not contain any exponential contribution. For positive *s*, the exponential contribution in the prefactor of erfcx(*s*) cancels the 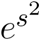 factor in the integrand. For negative *s*, the integrand simplifies even further to 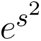 (1 + erf(–*s*)) = erfcx(*s*) using erf(–*s*) = – erf(*s*). In addition to erfcx(*s*), we employ the Dawson function

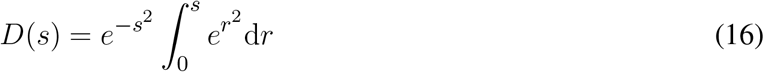

to solve some of the integrals analytically. The Dawson function has a power law tail, *D*(*s*) ~ 1/(2*s*); hence, it also does not carry an exponential contribution. Both erfcx(*s*) and the Dawson function are fully vectorized in `SciPy` (Virtanen et al., 2020).

Any remaining integrals are solved using Gauss–Legendre quadrature (Press et al., 2007). By construction, Gauss–Legendre quadrature of order *k* solves integrals of polynomials up to degree *k* on the interval [–1,1] exactly. Thus, it gives very good results if the integrand is well approximated by a polynomial of degree *k*. The quadrature rule itself is

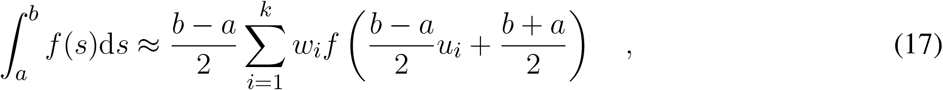

where the *u_i_* are the roots of the Legendre polynomial of order *k* and the *w_i_* are appropriate weights such that a polynomial of degree *k* is integrated exactly. We use a fixed order quadrature for which Eq. (17) is straightforward to vectorize to multiple *a* and *b*. We determine the order of the quadrature iteratively by comparison with an adaptive quadrature rule; usually, a small order *k* = *O*(10) already yields very good results for an erfcx(*s*) integrand.

#### Inhibitory Regime

First, we consider the case where lower and upper bound of the integral are positive, 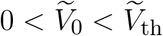. This corresponds to strongly inhibitory mean input. Expressing the integrand in terms of erfcx(*s*) and using the Dawson function, we get

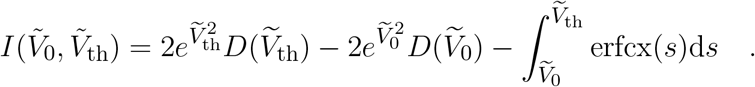

The remaining integral is evaluated using Gauss–Legendre quadrature, Eq. (17). We extract the leading contribution 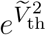 from the denominator in Eq. (2) and arrive at

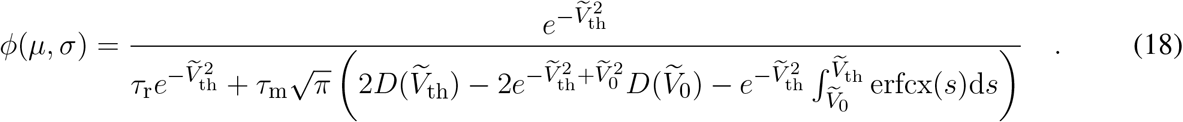

Extracting 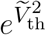 from the denominator reduces the latter to 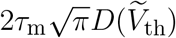 and exponentially small correction terms (remember 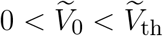 because *V*_0_ < *V*_th_), thereby preventing overflow.

#### Excitatory Regime

Second, we consider the case where lower and upper bound of the integral are negative, 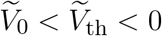. This corresponds to strongly excitatory mean input. In this regime, we change variables *s* → – *s* to make the domain of integration positive. Using erf(–*s*) = –erf(*s*) as well as erfcx(*s*), we get

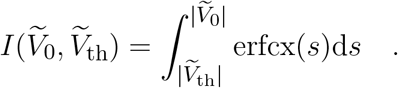

Thus, we evaluate Eq. (2) as

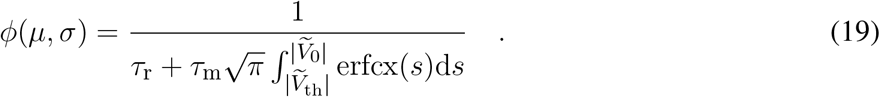

In particular, there is no exponential contribution involved in this regime.

#### Intermediate Regime

Last, we consider the remaining case 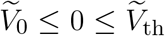. We split the integral at zero and use the previous steps for the respective parts to get

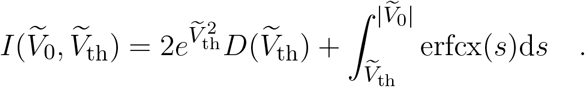

Note that the sign of the second integral depends on whether 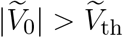 (+) or not (–). Again, we extract the leading contribution 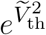 from the denominator in Eq. (2) and arrive at

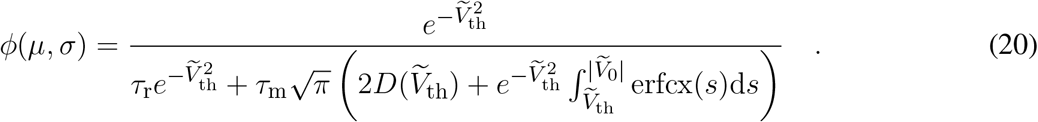

As before, extracting 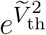 from the denominator prevents overflow.

#### Deterministic Limit

The deterministic limit *σ* → 0 corresponds to 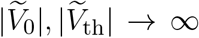 for both Eq. (3) and Eq. (7). In the inhibitory and the intermediate regime, we see immediately that *ϕ*(*μ*, *σ* → 0) → 0 due to the dominant contribution 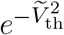. In the excitatory regime, we use the asymptotics 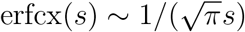 to get

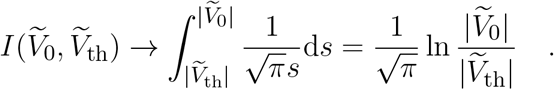

Inserting this into Eq. (2) yields

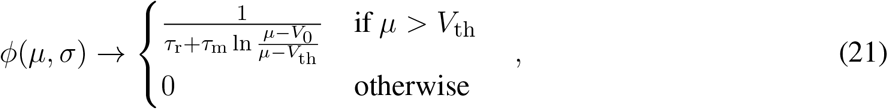

which is the firing rate of a leaky integrate-and-fire neuron driven by a constant input (Gerstner et al., 2014). Thus, this implementation also tolerates the deterministic limit of a very small noise intensity *σ*.

### 5.2 Microcircuit Parameters

**Table 1.** Parameter adaptions used here are introduced by Bos et al. (2016) compared to original micro-circuit model. *K_ij_* denotes the in-degrees from population *j* to population *i*. The delays in the simulated networks were drawn from a truncated Gaussian distribution with the given mean and standard deviation. The mean-field approximation of the microcircuit by Potjans and Diesmann (2014) assumes the delay to be fixed at the mean value, which is specified in the toolbox by setting the parameter `delay_dist` to `none`.

| Symbol | Value<br>Potjans and Diesmann (2014) | Value<br>Bos et al. (2016) | Description |
| --- | --- | --- | --- |
| $K_{4E,4I}$ | 795 | 675 | In-degree from 4I to 4E |
| $K_{4E,ext}$ | 2100 | 1780 | External in-degree to 4E |
| $D(\omega)$ | none | truncated Gaussian | Delay distribution |
| $d_e \pm \delta d_e$ | $1.5 \pm 0.75$ ms | $1.5 \pm 1.5$ ms | Mean and standard deviation of excitatory delay |
| $d_i \pm \delta d_i$ | $0.75 \pm 0.375$ ms | $0.75 \pm 0.75$ ms | Mean and standard deviation of inhibitory delay |

## Footnotes

1 https://github.com/INM-6/nnmt

2 http://alleninstitute.github.io/dipde

3 https://nnmt.readthedocs.io/

4 https://nnmt.readthedocs.io/

## Notes

### Competing Interest Statement

The authors have declared no competing interest.

